# Auditory midbrain encodes training-induced plasticity in sound localization behavior

**DOI:** 10.1101/2025.09.22.677707

**Authors:** Ana Sanchez Jimenez, Victoria M. Bajo, Ben D. B. Willmore, Andrew J. King, Fernando R. Nodal

## Abstract

The ability of the brain to learn from experience and to compensate for changes in sensory inputs is usually associated with plasticity in the cerebral cortex. Descending corticofugal pathways have been implicated in learning but there is limited evidence that subcortical processing is shaped by training. We show that sound-source location can be accurately decoded from neuronal populations recorded in the inferior colliculus of one or both hemispheres while ferrets perform a localization task. Furthermore, changes in neural decoding performance matched improvements in localization accuracy of individual animals when ferrets were trained to adapt to abnormal spatial cues resulting from reversible occlusion of one ear. These findings demonstrate that the activity patterns of populations of neurons in the inferior colliculus can account for sound localization behavior in different hearing contexts, and that training-dependent plasticity in the auditory midbrain may support spatial learning following monaural hearing loss.

## Introduction

An ability to localize sound sources accurately and rapidly is a crucial feature of many predator-prey relationships and makes an important contribution to human auditory perception in listening environments that often include multiple sources, echoes and reverberation^1^. This is a complex task since sound source locations must be computed from binaural and monaural spatial cues generated by the geometry of the head and external ears. Neural sensitivity to interaural time differences (ITDs), interaural level differences (ILDs) and monaural spectral cues arises in separate auditory brainstem nuclei^1,2^. The inferior colliculus (IC) is the first site of integration of these frequency-dependent spatial cues^3,4^, a critical step in sound localization, and is an obligate relay for most of the auditory information transmitted via the thalamus to the auditory cortex^5–7^, as well as a major source of auditory input to the superior colliculus, where a map of auditory space is found^8^.

The neural circuits responsible for representing sound source location are shaped throughout life by experience^9^. This plasticity is essential for calibrating the circuits as the head and external ears grow during development. Importantly from a clinical perspective, plasticity also provides a basis for adapting to altered spatial cues resulting from hearing disorders^10,11^. Indeed, the adult brain possesses a remarkable capacity to learn to utilize experimentally altered spatial cues^12^, which, like other forms of auditory perceptual learning^13–17^ is thought to involve the auditory cortex^18–20^. However, there is growing evidence that while the primary auditory cortex (A1) plays a critical role in learning, the resulting neural plasticity may be expressed and consolidated in downstream circuits^14,18^. Thus, the capacity of adult ferrets to relearn to localize sound accurately after altering the relationship between auditory spatial cues and sound source location is similarly impaired by silencing A1 during training^18^ and by selectively eliminating layer 5 neurons that project to the IC^21^, highlighting the need to investigate plasticity at subcortical levels too.

The broad and predominantly contralateral tuning of neurons in the mammalian IC led to the notion that the azimuthal location of sound sources is encoded by differences in overall activity between each hemisphere^2,22^. Other work has instead argued that decoding sound azimuth from the activity patterns across populations of IC neurons provides a better fit to the localization abilities of the species used^23–26^. None of these studies measured IC activity in behaving animals, however, which is essential for determining the functional relevance of these neural coding strategies.

We chronically recorded neural activity in the IC of ferrets while they localized sounds in the azimuthal plane, both under normal hearing conditions and when they experienced abnormal spatial cues resulting from a reversible conductive hearing loss in one ear. Our findings provide new insights into the representation of auditory space in the midbrain and the extent to which the localization accuracy of individual animals can be accounted for by decoding activity from IC neurons in one or both hemispheres. We also show that the neural decoder performance changes across testing sessions to match localization accuracy during adaptation to monaural hearing loss, indicating an important role for the auditory midbrain in the neural plasticity that supports spatial learning in adults.

## Results

### Response types in the inferior colliculus of awake behaving ferrets

The aims of this study were to examine the relationship between the spatial response properties of neurons in the auditory midbrain and localization accuracy of the same animals, and whether those properties change following monaural occlusion in ways that could account for the training-induced adaptation in the animals’ behavior.

Ferrets were trained by positive reinforcement in an approach-to-target sound localization task (Fig. 1a). They performed almost perfectly at the longest stimulus durations (≥1000 ms) tested, with their accuracy declining at shorter durations, as previously reported^27^ (Fig. 1b). Once trained, a subset of 3 ferrets was chronically implanted with Neuropixels 1.0 probes in the IC (bilaterally n=2, left side n=1). Single and multi-unit activity was recorded on each probe while the ferrets performed the sound localization task, both under normal hearing conditions and after altering the spatial cues available by reversibly occluding one ear^28,29^.

The probes were positioned dorsoventrally in the IC (Fig. 1c, Extended Fig. 1a), spanning the whole central nucleus (CNIC) (approximately 2 mm in depth), with some of the most dorsal recording sites located in the overlying dorsal cortex (DCIC) (Fig. 1d, green bars). The ferrets were temporarily sedated to facilitate detailed characterization of frequency response areas along the probes. In keeping with previous work^30^, CNIC units were tuned for sound frequency, and their characteristic frequencies exhibited a dorsoventral progression from low to high (Fig. 1e, Extended Fig. 1b).

**Fig. 1:**
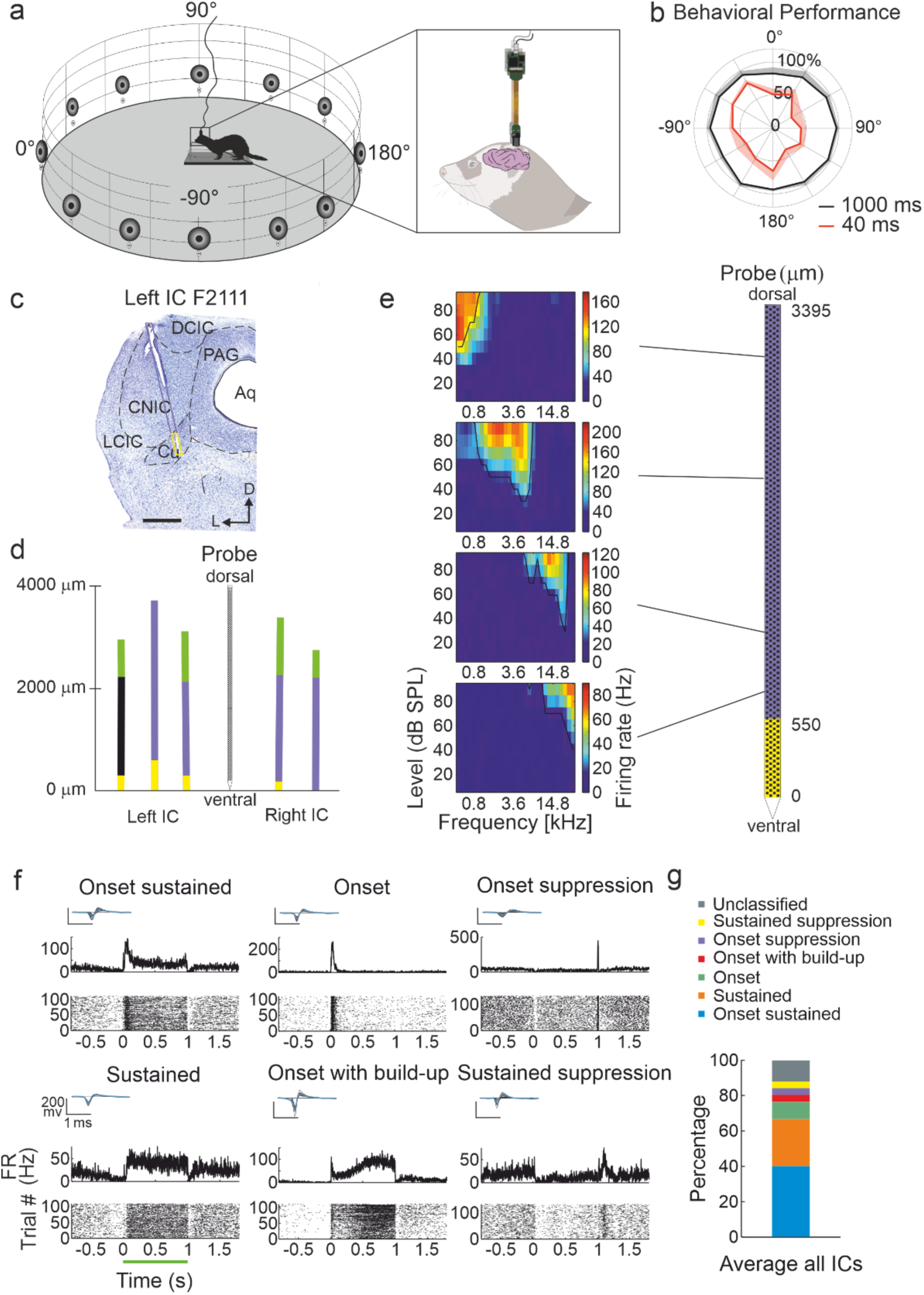
Recording setup and responses of IC neurons in awake ferrets. **a**, Schematic of the behavioral arena for the sound localization task while recording neural activity in the IC of ferrets. The animal stood on the central platform to trigger a burst of broadband noise from one of the 12 peripheral loudspeakers, and then had to approach the corresponding waterspout to receive a reward. **b**, Sound localization performance averaged across animals for two stimulus durations, 1000 ms in black and 40 ms in red. Percentage correct responses are plotted for different azimuthal angles. Negative locations correspond to positions to the left, and positive to the right of the location in front of the ferret (0°). **c**, Coronal section stained for Nissl substance at the level of the left IC in case F2111, with the chronically implanted Neuropixels probe superimposed on the electrode track and color coded according to its anatomical location. Aq, Aqueduct; DCIC, dorsal cortex of the IC; Cu, cuneiform nucleus; CNIC, central nucleus of the IC; LCIC, lateral cortex of the IC; D, dorsal; L, lateral. Scale bar, 1 mm. **d**, Anatomical location of each recording probe in different midbrain areas (n = 3, left side of the brain, n = 2, right side): green, DCIC; violet, CNIC; yellow, Cu; black, caudal pole of CNIC. **e**, Dorsoventral tonotopy illustrated by example frequency response areas (50-ms tones presented at the frequencies and levels shown) recorded at different depths along the recording probe in the left IC of case F2111, as shown in **c**. **f**, Peristimulus histograms and raster plots for example units illustrating the range of temporal firing patterns recorded in response to 1000-ms broadband noise bursts presented during the localization task. Insets represent the action potential waveform used to identify neural clusters. **g**, Proportion of each response type across all animals.

A variety of sound-evoked spiking patterns were observed in response to the broadband noise bursts used to measure spatial tuning during the localization task. Responses were classified into 6 main types based on spiking activity across 3 different windows during the sound presentation (Fig. 1f). Most of the units produced an excitatory response (80.42 ± 6.02%) and were further classified into ‘onset’ if the increase in firing rate was constrained to the first 50 ms of the sound, ‘onset sustained’ when adaptation followed the onset response, and ‘sustained’ and ‘onset with build-up’ when that was not the case. The two most common response types were ‘onset sustained’ (40.03 ± 9.27%) and ‘sustained’ (26.72 ± 5.78%) (Fig. 1g, Extended Fig. 2). A minority of units showed inhibitory responses to sound (11.47 ± 7.38 %), characterized by a decrease in firing rate following the stimulus onset relative to preceding activity. This suppression was either restricted to the onset of the sound (‘onset suppression’) or continued throughout the sound presentation (‘sustained suppression’). In suppressed units, an excitatory offset response was typically observed, probably due to a post-inhibitory rebound.

### Spatial tuning of IC neurons in behaving ferrets with normal binaural hearing

The activity of IC neurons varied with the location of the sounds presented during the task. Their spatial preferences, estimated by the vector sum of their firing rates at different azimuths (the centroid), varied widely but generally showed a clear preference for stimuli presented in the contralateral hemifield (Fig. 2a,b). This contralateral preference was found in both the left and right IC across all animals (median centroid left IC: 77.21°, right IC: -94.67°). However, in each case, a minority of units exhibited an ipsilateral preference (Fig. 2b). Reflecting the generally broad response-azimuth functions of the units (Fig. 2a), the equivalent rectangular receptive field (ERRF) widths ranged from 50°-260°, with a mean >150° (Fig. 2c).

**Fig. 2:**
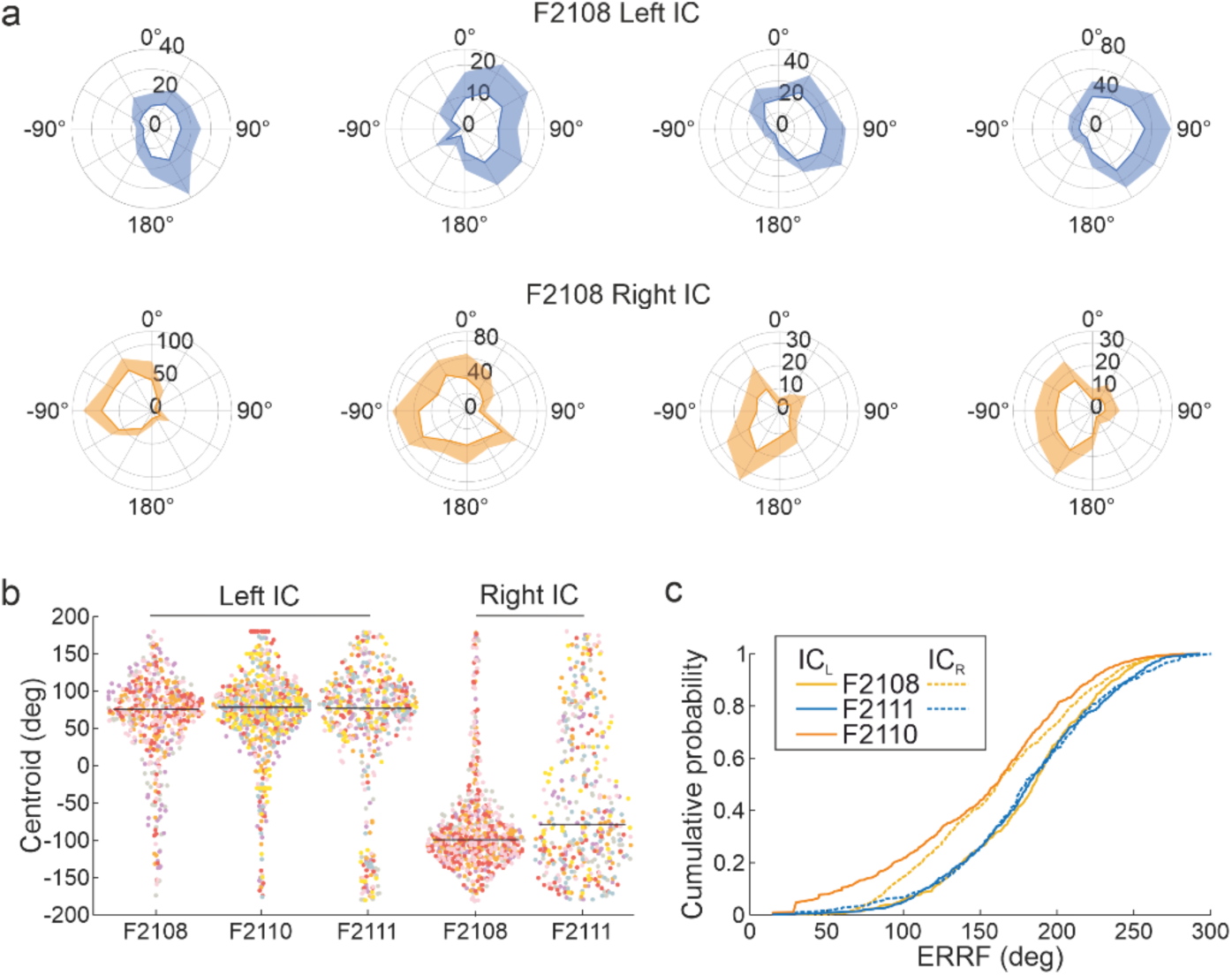
Spatial response properties of IC neurons. **a**, Polar plots illustrating the broad spatial tuning of example units recorded in the left IC (top row) and right IC (bottom row; case F2108) during the sound localization task. Lines indicate the mean firing rate (spikes/s) and the shaded areas are the standard deviation. **b**, Distribution of the preferred azimuthal direction (centroids) for units (dots) recorded over several sessions (different colors) for each animal and each IC. Horizontal lines depict the median centroid. Most units had a contralateral preferred direction. **c**, Cumulative distribution of the equivalent rectangular receptive field (ERRF) width for the units shown in **b**.

### A linear model can decode sound location from the IC

We used a population decoding analysis to explore to what extent sound location could be decoded from the information carried by the responses of IC neurons recorded during the localization task. A linear pattern model was trained for each behavioral session for each animal using the activity recorded bilaterally from the IC. Because the proportion and type (left-right vs back-front) of localization errors varies with target location^31^, we hypothesized that the neural representations of left-right and back-front location are linearly separable. The decoder therefore estimated the target azimuth in degrees from independent projections onto the left-right (cosine of angular location) and back-front (sine of angular location) axes. Additionally, the model provided information about the weighting of each unit’s response in each time bin for the decoding of each axis.

In the left-right axis, the absolute values of the weights indicated that IC neurons were most informative about locations in the opposite hemifield (Fig. 3a, Extended Fig. 3a). This is consistent with their overall contralateral preference. The highest absolute weights were found in the first 200 ms of spiking activity after sound onset, corresponding to the time window of maximum evoked activity for most of the units with either onset or onset-sustained responses. Conversely, the absolute weights in the back-front axis were more homogeneous throughout the stimulus (Fig. 3b, Extended Fig. 3b), and the decoding error was larger than for the left-right axis (Fig. 3c, Extended Fig. 3c). The early peak in neural weights and the superior decoding performance of the model in the left-right dimension supports previous observations that the accuracy of the initial head orienting response made by ferrets in this task is driven by the onset of the target sound, and that left-right errors are rare even at short sound durations^31^.

Further evidence that IC activity is sufficient to account for auditory localization accuracy in ferrets is provided by the close match between the decoded and actual azimuth locations (Fig. 3d-f) and the near-perfect behavioral performance of the animals for the 1000-ms noise bursts presented as target sounds (Fig. 1b). Because ferrets orient toward the sound source with a latency of ∼200 ms^31,32^, these stimuli are long enough to potentially provide dynamic spatial cues that may have influenced neural decoding performance. As a control for this, we also trained the model with spiking activity restricted to a 200-ms window after sound onset. With this shorter response window, the model was still able to decode target location with an accuracy comparable to that observed with the full trial duration (Extended Fig. 3d-f), further indicating that most spatial information is conveyed by the onset responses and ruling out any major contribution of dynamic spatial cues to the performance of the neural decoder. This again fits with the behavioral data; while the approach-to-target performance declines as the duration of the target sound is reduced, the accuracy of the initial head orienting responses is much less dependent on stimulus duration^31^.

**Fig. 3:**
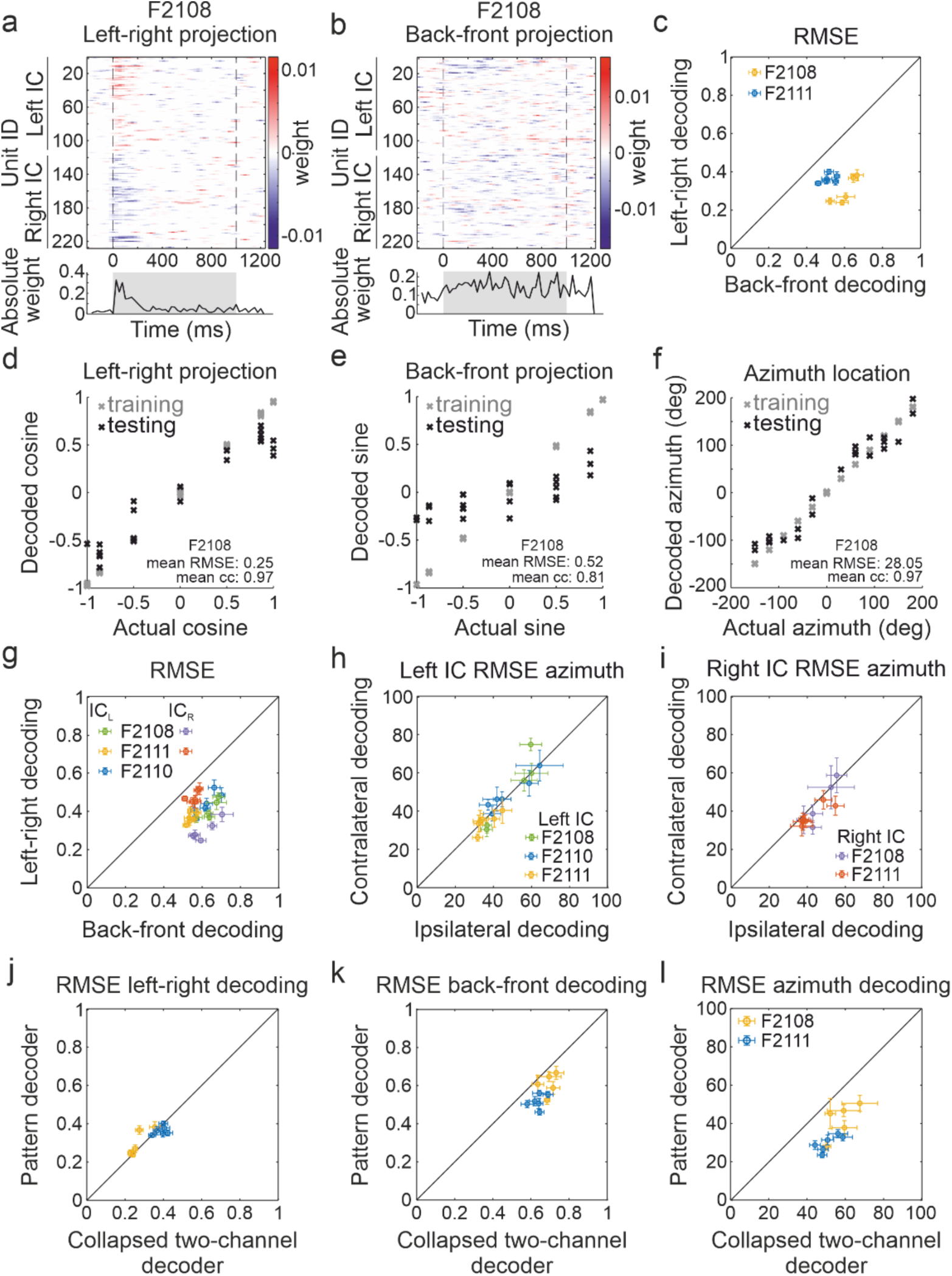
A simple linear pattern model captures spatial information encoded in the IC. The 12 target locations used in the sound localization task were decomposed into their left-right and back-front projections, which were then decoded from the sound-driven responses using a linear population-pattern model. For each session, the number of units recorded in each IC was balanced, and the model was independently fitted multiple times (*k*-folds). For each fold, 12 different trials were selected for testing the model and training was performed with the remaining trials from that session. **a**, **b**, Temporal distribution (relative to sound presentation) of the unit weights for an example fold of the model (case F2108) for the left-right projection (**a**) and the back-front projection (**b**). Units are ordered according to their IC location in each hemisphere, and the color indicates how informative each unit is about the decoded variable (blue: left hemifield (**a**), back (i.e., posterior hemifield, **b**); red: right hemifield (**a**), front (anterior hemifield, **b**)). Panels below show the sum of absolute weights from all units, which peaked after sound onset for decoding the left-right axis. **c**, Root mean squared errors (RMSEs) for decoding the left-right vs the back-front axis for animals with bilateral IC recordings. **d**, **e**, **f**, Distribution of values for the training trials (gray) and testing trials (black) for decoding the left-right projection (cosine), back-front projection (sine), and azimuthal location (degrees), respectively, for an example fold. Overall performance across folds is expressed as the mean RMSE and mean correlation coefficient. **g**, **h**, **i**, Decoding performance based on either the left or the right IC only. **g**, RMSEs for decoding the left-right vs the back-front axis for one IC. **h**, **i**, Comparison of RMSEs for decoding contralateral vs ipsilateral locations for models using only left (**h**) or right IC units (**i**). **j**, **k**, **l**, Comparison of RMSEs between the population-pattern and collapsed two-channel decoders for the left-right axis (**j**), back-front axis (**k**) and azimuthal location (**l**).

Unilateral lesions of the IC result in predominantly contralateral deficits in sound localization^33–36^, implying that each side of the midbrain represents azimuthal locations in the opposite hemifield. Because units in the IC exhibit a mix of predominantly contralateral but also some ipsilateral spatial preferences, it is possible that sound location may be encoded by the relative activity of these two populations within each side of the brain, as suggested previously for the auditory cortex^37,38^. We therefore trained the model to classify the azimuthal angle of the stimulus using the same number of neurons recorded from one side of the brain only. As with the bilateral IC model, decoder performance using one IC was better in the left-right axis than the back-front axis (Fig. 3g, Extended Fig. 3g). Root mean square errors (RMSE) between the actual and predicted locations were, however, slightly larger for one IC than when data from both sides were included (two-way repeated measures ANOVA: effect of brain location (both sides, left IC, right IC), F_1,59_ = 3.37, p = 0.072; interaction between brain location and decoding dimension, F_1,59_ = 33.05, p <0.0001; pairwise comparison of left-right decoding for both sides vs left IC only, p=0.022) (Fig. 3c,g). The poorer performance with a single IC was not due to an asymmetry in decoding accuracy between contralateral and ipsilateral locations (Fig. 3h,i, Extended Fig. 3).

The superior decoding performance obtained by combining information from both sides of the brain is potentially consistent with the hypothesis that spatial locations are encoded by the relative activity of two broad, hemispheric channels in the left and right IC^22^. To further examine this, we built a variant of the population decoder by training the model with activity in each IC collapsed into a single ‘channel’ by averaging the spiking activity across all recorded units to preserve their temporal firing patterns (Fig. 3j-l, Extended Fig. 3j-l). Decoding performance for sound location in the left-right axis was very similar in the two models (Fig. 3j, Extended Fig. 3j). However, for back-front (Fig. 3k, Extended Fig. 3k) and azimuth decoding (Fig. 3l, Extended Fig. 3l), the pattern decoder consistently outperformed the two-opponent channel model. The pattern decoder was, therefore, able to extract more information from the population spiking activity than just the comparison of the average responses between each IC.

### Monaural occlusion changes the spatial tuning of IC neurons

To further understand how spatial information is encoded in the IC, we altered the available spatial cues by plugging the left ear and examined whether the resulting changes in their spatial response properties could account for the effects of this unilateral conductive hearing loss on the animals’ localization behavior (Fig. 4). The immediate effect of the earplug, which attenuates (by 15-45 dB) and delays (by 110 µs) the acoustical input^29^, was to change most of the sound-evoked excitatory responses in the contralateral IC to inhibition, as shown by a significant reduction in the normalized firing rate compared to the pre-plug recordings (Wilcoxon signed-rank test, n = 2 ferrets, W = 789430, p <0.0001) (Fig. 4c). By contrast, in the left IC, plugging the ipsilateral left ear had no effect on the firing rate of the neurons (Wilcoxon signed-rank test, n = 3 ferrets, W = 520840, p = 0.23) (Fig. 4a). This can be explained by the binaural imbalance caused by the earplug on sound-driven responses in the auditory brainstem and the laterality of the projections from these nuclei to the IC (Fig. 4b).

The behavioral responses of these animals showed a marked reduction in localization accuracy when the left ear was first plugged (Fig. 4d, Extended Fig. 4a,h), with a rightward bias toward the side of the open right ear^27,32^. This was accompanied by a loss of the contralateral dominance in the ipsilateral left IC (Wilcoxon signed-rank test, n = 3 ferrets, W = 82620, p <0.0001) (Fig. 4e, Extended Fig. 4), with the average centroid shifted towards the midline (0°) and an increase in ERRF width (Wilcoxon signed-rank test, n = 3 ferrets, W = 36273, p <0.0001 (Fig. 4e, Extended Fig. 4). The degraded spatial tuning is most likely explained by a loss of inhibitory influence from the occluded ear on binaural processing. Together with the dramatic suppression of activity in the right IC, contralateral to the occluded ear, (Fig. 4f, Extended Fig. 4) these changes disrupted the representation of sound azimuth in the midbrain, with units in both hemispheres showing more omnidirectional responses (Fig. 4g-j, Extended Fig. 4).

**Fig. 4:**
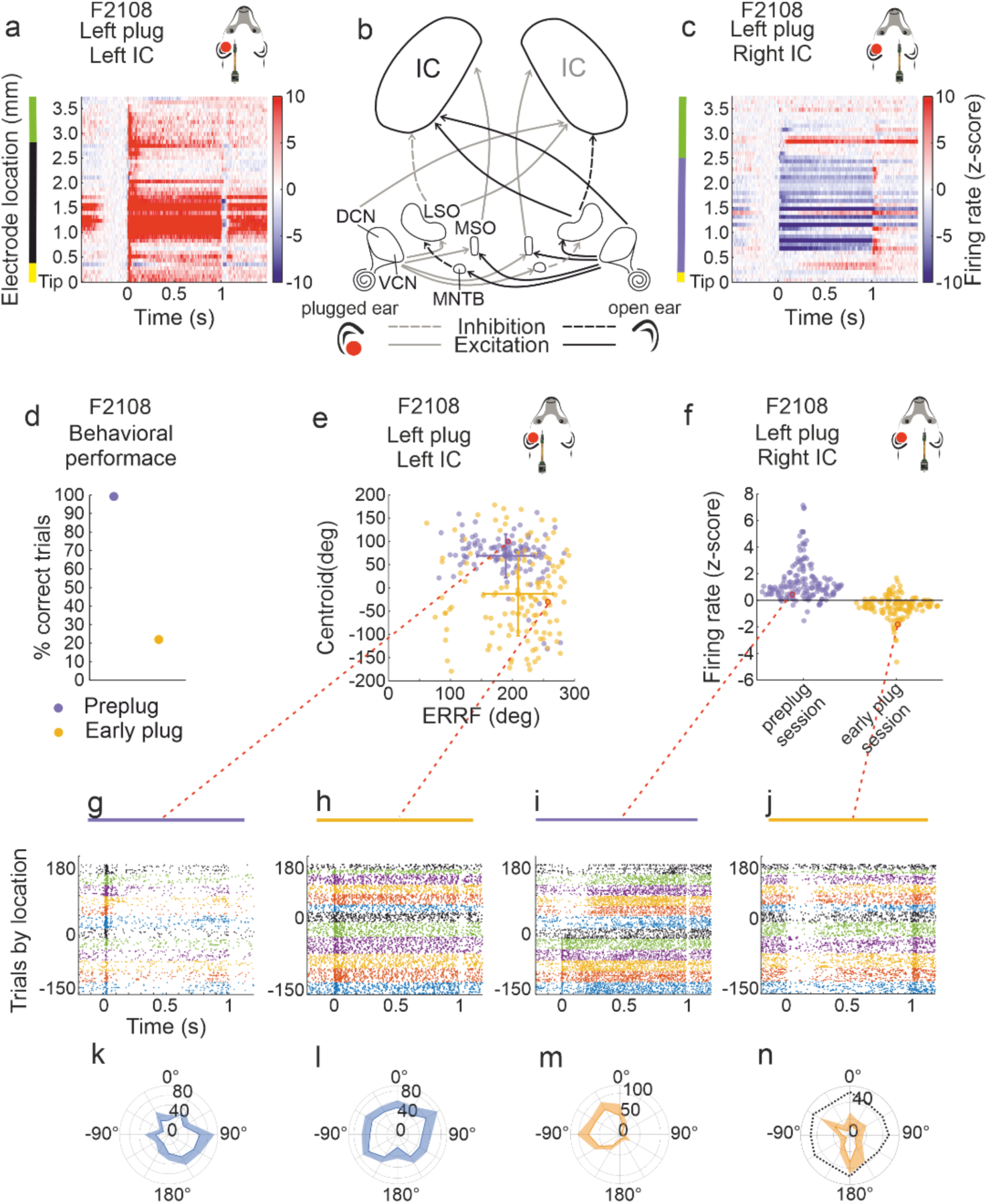
Effects of monaural hearing loss on IC activity and spatial response properties. **a**, **b**, **c**, Insertion of an earplug in the left ear altered the balance of sound-driven activity between the left and right IC. PSTHs organized by depth along the recording probe show predominantly excitatory responses in the ipsilateral left IC (**a**) and inhibitory responses in the contralateral right IC (**c**), which are predicted by the binaural imbalance in the auditory brainstem resulting from the reduced input from the plugged ear (**b**). **d**, Initial reduction in localization accuracy caused by monaural hearing loss in case F2108. **e**, Changes in spatial response properties (centroid and ERRF width) of all units recorded in the left IC of F2108 following earplug insertion. **f**, Suppression of sound-driven activity in the right IC of F2108 following earplug insertion. Raster plots of example units in the left IC (**g**, **h**) and right IC (**i**, **j**) before and after earplug insertion. **k**-**n**, Azimuth response profiles of these same units. The unit in **j** and **n** was inhibited by sound and its baseline activity is shown by the dotted line in **n**.

### Plasticity of spatial tuning in IC neurons mirrors perceptual learning

Following the initial drop in behavioral performance when one ear was occluded, subsequent sound-localization training with the earplug still in place resulted in a gradual improvement in accuracy across training sessions and spatial locations in all animals, including those with cranial implants (Fig. 5a). To explore whether behavioral adaptation to unilateral conductive hearing loss was associated with plasticity in the spatial response properties of IC neurons, we trained the linear pattern decoder to classify target location using bilateral neural activity from each session separately.

The initial effect of the earplug was to degrade IC decoding in both the left-right and back-front dimensions, producing large mismatches between the decoded and actual azimuthal locations (illustrated for F2108 in Fig. 5b-d). We then examined the decoding performance of the linear pattern model at different stages of adaptation to unilateral conductive hearing loss, which revealed a significant difference in both left-right RMSE (n = 2, F_1,106_ = 6.121, p = 0.0149) and azimuth decoding RMSE (n = 2, F_1,106_ = 5.659, p = 0.0192) between the first and second halves of the sessions with the earplug in place.

Across these sessions, the azimuth decoding errors estimated from the bilateral IC responses aligned closely with the errors in localization behavior (Fig. 5e,f). This was the case irrespective of the degree of behavioral adaptation to unilateral conductive hearing loss. Thus, in F2108, a clear improvement in localization accuracy was observed with training, as reflected by a significant reduction in RMSE across sessions (linear regression R^2^ = 0.59, slope = -1.36, p = 1.75e-06), which was accompanied by a comparable reduction in azimuth decoding RMSE estimated from IC activity over the full stimulus duration (R^2^ = 0.49, slope = -1.02, p = 3.14e-05) (Fig. 5e). In contrast, F2111 showed little change over the period of monaural occlusion in either behavioral (R^2^ = 0.11, slope = -0.39, p = 0.0513) or neural decoding (R^2^ = -0.01, slope = -0.22, p = 0.377) errors (Fig. 5f). A significant correlation was nonetheless found for this ferret between behavioral and decoding errors (R^2^ = 0.125, slope = 0.33, p = 0.0367), indicating that population activity in the IC was still informative about the animal’s behavior.

When normal binaural cues are available, IC activity in the period following sound onset is most informative about the target location, as shown by greater absolute weights in the decoding kernels for the left-right axis (Fig. 3a). When the ear was first occluded, however, the weights no longer peaked during the first 200 ms of the sound presentation (Fig. 5g). The subsequent improvement in azimuth decoding was due primarily to a reduction in left-right decoding errors (F2108, R^2^ = 0.57, slope = -0.006, p = 3.42e-06), with a clear increase in the post-stimulus onset weights between the first three (Early plug) and last three (Late plug) training sessions (Fig. 5g-i). This recovery in left-right decoding accuracy was particularly pronounced in the right IC (Fig. 5h,i).

In the case of F2108, the animal showing the clearest improvement with training in both behavioral and neural decoding performance (Fig. 5e), the firing rate z-score in the right IC showed a significant gradual increase across the sessions in which the ear was plugged (Extended Fig. 5a). This was accompanied by a partial recovery of the contralateral preference of these neurons, as shown by the emergence of a difference in their weighted average firing rate in response to sounds presented in the left versus the right hemifield (Extended Fig. 5b,c) and a shift in the population average centroid towards the contralateral side (Extended Fig. 5c,d). These changes are likely to account for the increase in left-right decoding weights in the right IC and be part of the compensatory process that underpins behavioral adaptation to unilateral conductive hearing loss.

Training the linear pattern decoder with the spiking activity recorded in both ICs during the first 200 ms after sound presentation revealed same trend as for the full stimulus duration in both ferrets F2108 (RMSE for 200-ms window following stimulus onset, R^2^ = 0.51, slope = -0.94, p = 1.67e-05) and F2111 (R^2^ = -0.04, slope = -0.03, p = 0.907) (Extended Fig. 6). This again indicates that onset activity in the IC is the most informative part of the response and able to account for the animals’ localization behavior, minimizing the possible contribution of dynamic spatial cues once the ferret starts to move its head to the improvement in neural decoding performance.

**Fig. 5:**
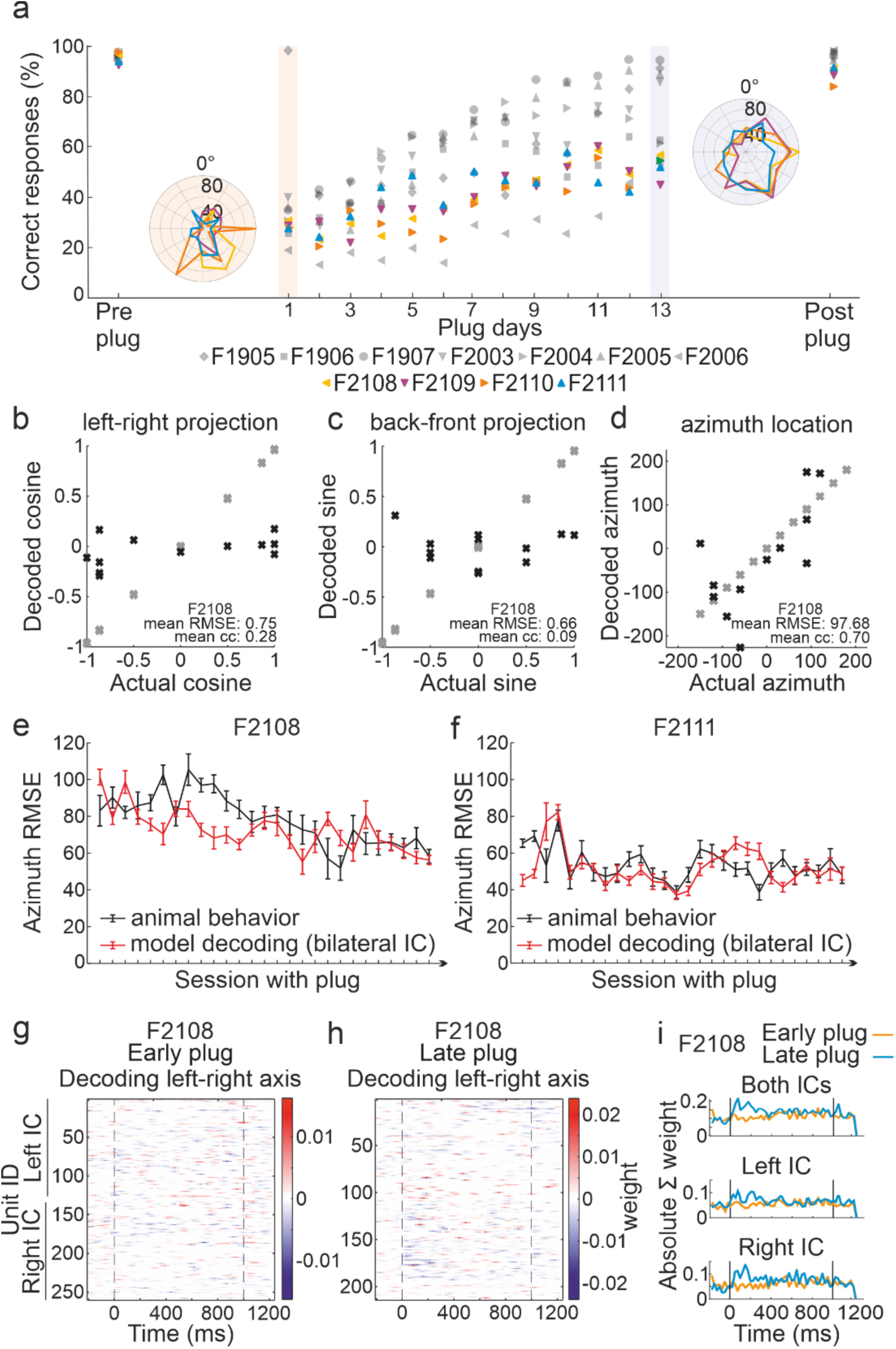
Training-dependent adaptation in sound localization behavior is captured by decoding spiking activity in the inferior colliculus. **a**, Percentage correct scores averaged across the 12 loudspeaker locations are plotted for different animals before, during and after occlusion of the left ear; colored symbols indicate the animals with Neuropixels probes implanted in the IC. The earplug was in place for up to 20 days (days 1–13 only are shown). Polar plots show the mean score for the implanted ferrets at different target locations in the first few days (left) and toward the end of the earplugging period (right). **b**-**d**, Distribution of values for the training trials (gray) and testing trials (black) for decoding the left-right axis (**b**), back-front axis (**c**), and azimuthal location (**d**) from IC activity for an example fold at the start of the earplugging run in case F2108. Overall performance across folds is expressed as the mean RMSE and mean correlation coefficient. **e**, **f**, Behavioral (black) and neural decoding (red) errors across the earplugging sessions for the animals with bilateral IC recordings (**e**, F2108; **f**, F2111). **g**, **h**, Temporal distribution (relative to sound presentation) of the unit weights for an example fold of the population-pattern model (F2108) for decoding the left-right axis in an early (**g**) and late (**h**) earplugging session. Units are ordered according to their IC location in each hemisphere, and the color indicates how informative each unit is about the decoded variable (blue: left hemifield; red: right hemifield). **i**, Sum of absolute decoding weights for all units in the first three (early plug) and last three (late plug) sessions based on recordings from both sides of the brain (top), the left IC only (middle), and the right IC only (bottom) in case F2108.

To determine whether adaptation of localization requires activity in both ICs, as predicted by the opponent hemispheric model of azimuth coding, we trained the linear pattern decoder to classify stimuli along the left-right axis using unilateral IC activity only (left IC, ipsilateral to the earplug, n = 3) from the sessions in which the animals experienced monaural hearing loss. In all ferrets, a significant correlation was found between behavioral and neural decoding performance for the 1000-ms noise bursts used in these experiments (Fig. 6) and this remained the case in two of the animals (F2108 and F2111) when left IC decoding activity was restricted to the first 200 ms after sound onset (Extended Fig. 7).

These results therefore suggest that adaptive changes in spatial processing in the left IC, contralateral to the non-occluded ear, can support the recovery in sound localization accuracy. Indeed, in the ferrets in which bilateral recordings were made, no difference was found in azimuth (F1,106 = 0.734, p = 0.3935) or left-right (F1,106 = 0.274, p = 0.6018) decoding errors when the models were trained using spiking activity recorded from the left IC versus both hemispheres.

Finally, we compared the performance of the population-pattern and two-opponent channel models over the sessions in which ferrets were trained on the localization task with one ear occluded. In both ferrets with bilateral IC recordings, the pattern decoder clearly outperformed the two-opponent channel model (azimuth RMSE, F1,106 = 62.918, p < 0.001; left-right RMSE, F1,106 = 56.606, p < 0.001) (Fig. 7).

**Fig. 6:**
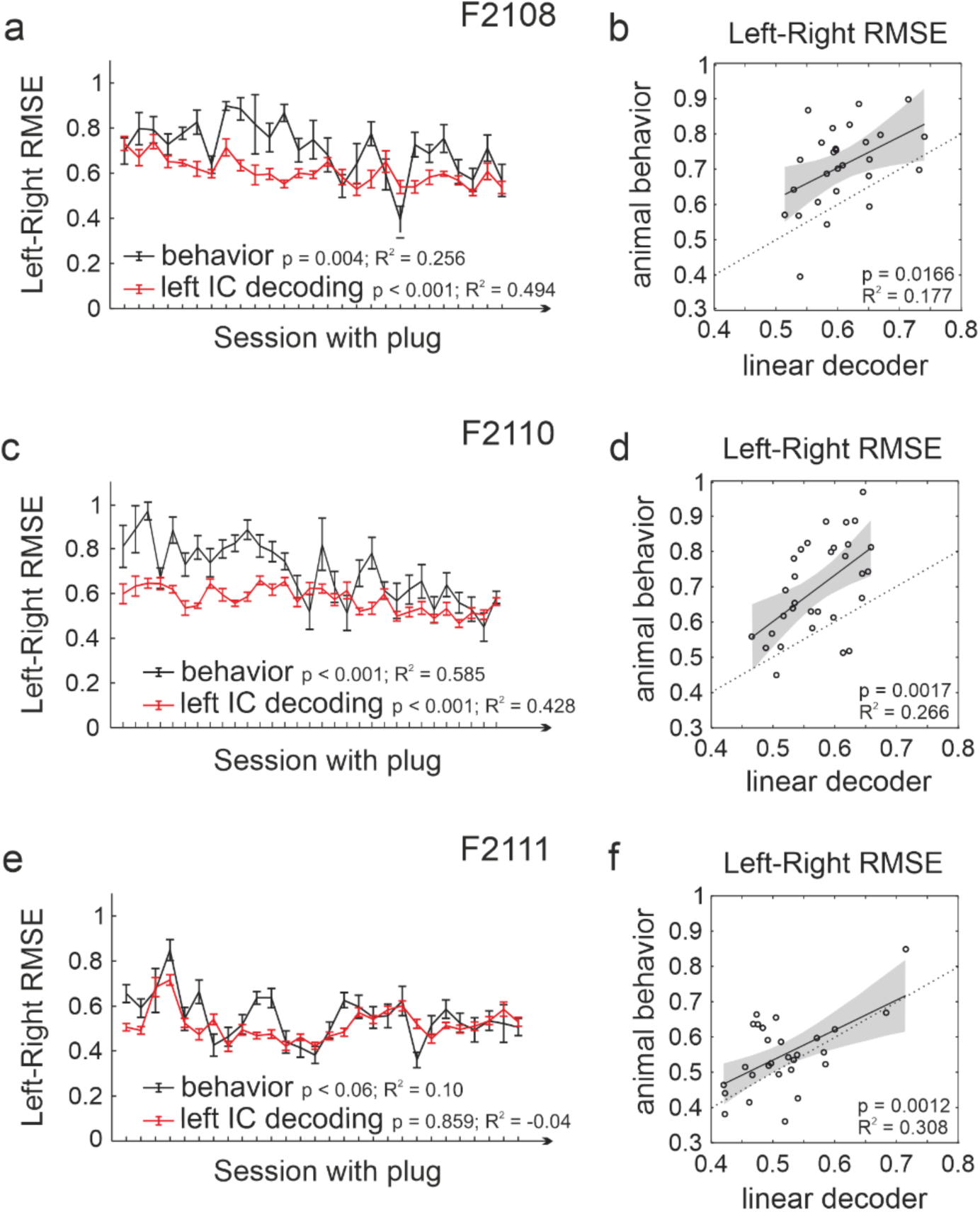
Decoding localization performance from neural activity in one IC during ear plugging. **a**, **b**, Comparison of behavioral and left-right population-pattern decoding errors during monaural hearing loss using left IC activity only in case F2108. **c**, **d**, Same for F2110. **e**, **f**, Same for F2111. **a**, **c**, **e**, Covariation of behavioral (black) and IC decoder (red) RMSEs across all earplugged sessions. **b**, **d**, **f**, Linear regression between the behavioral and decoder RMSEs in each session and animal.

**Fig. 7:**
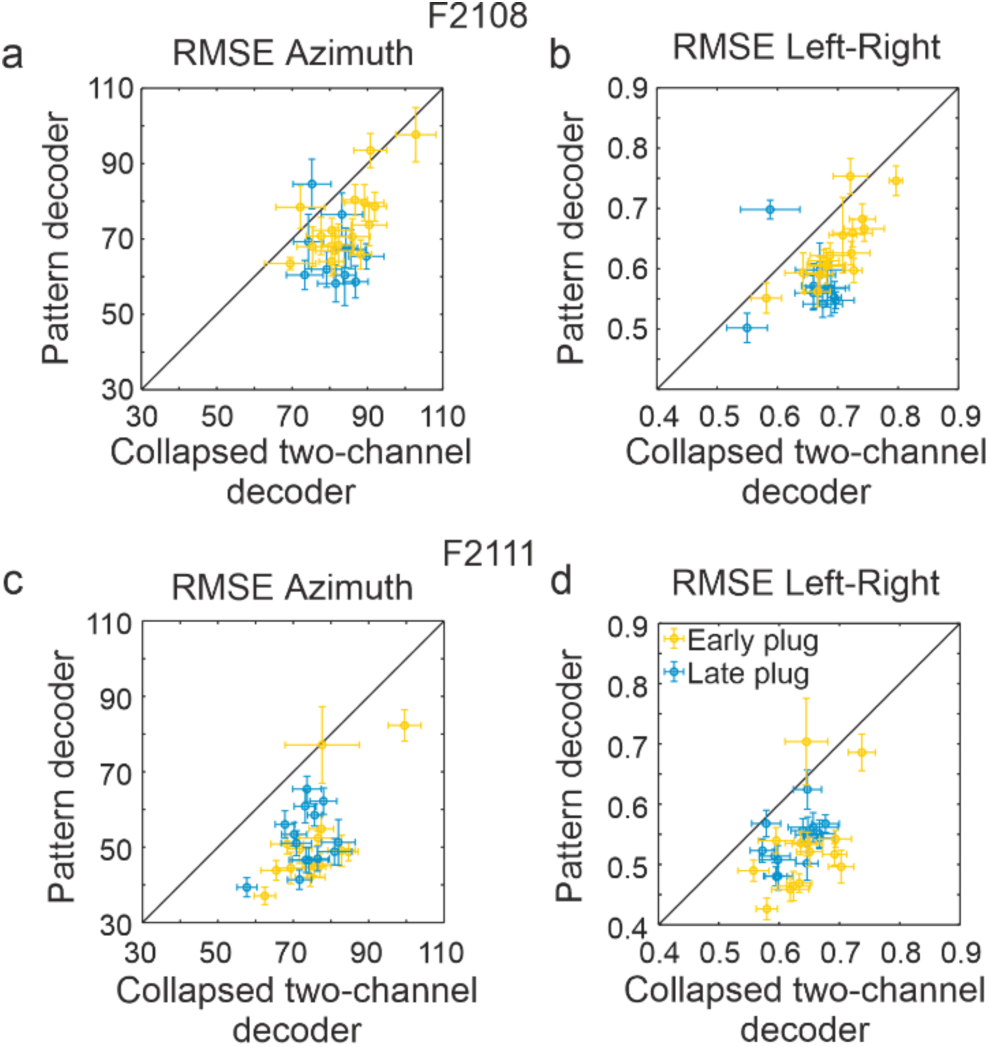
Comparison between population-pattern and collapsed two-opponent channel decoders during adaptation to monaural hearing loss. **a**-**d**, Comparison of decoding RMSE in the cases with bilateral IC recordings (F2108 **a**, **b**; F2111, **c**, **d**) for the population-pattern and two-opponent channel models over the sessions in which ferrets were trained on the localization task with one ear occluded. In both ferrets with bilateral IC recordings, the pattern decoder clearly outperformed the two-opponent channel model for both azimuth location (**a**, **c**) and left-right axis (**b**, **d**), independently of the amount of time the animals had one ear occluded (yellow, early plug sessions; blue, late plug sessions).

### Parallel recovery of behavioral and neural responses following earplug removal

After training the ferrets with one ear occluded for approximately 20 days, the earplug was removed to allow them to experience normal binaural inputs again. The accuracy of the post-plug approach-to-target responses immediately increased toward pre-plug levels (Fig. 8a). Again, these behavioral changes were closely mirrored by the responses recorded in the IC, which showed a comparable decrease in azimuth decoding errors (Fig. 8b). Furthermore, in both hemispheres, the early peak in the absolute weights in the decoding model (Fig. 8c) and the contralateral dominance of the centroids of the azimuth response profiles (Fig. 8d) were largely restored. These results further show that IC spatial sensitivity changes in parallel with the plasticity in sound localization behavior.

**Fig. 8:**
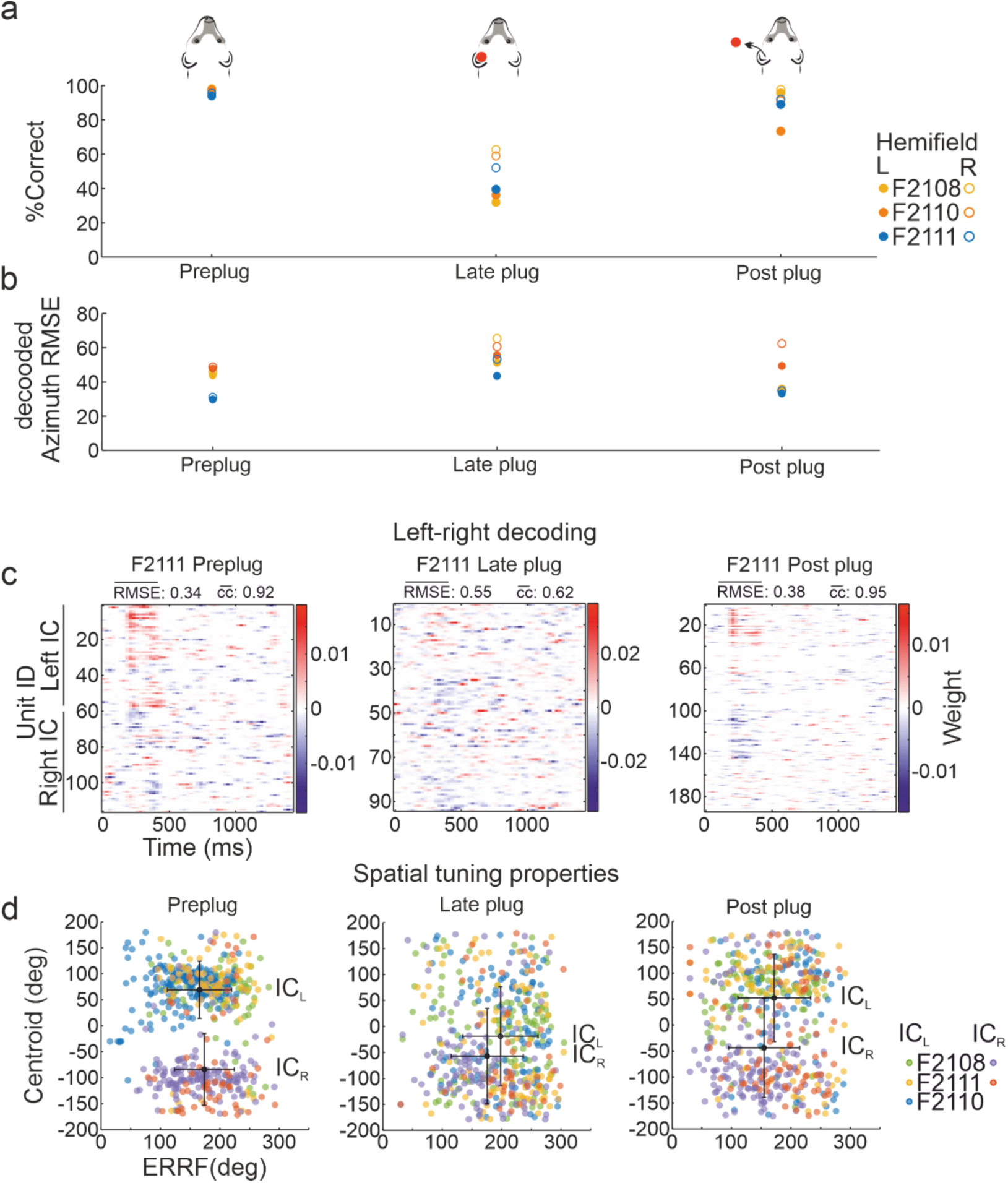
Sound localization accuracy and IC spatial properties recover in parallel following earplug removal. **a**, After ∼20 days of training with the left ear occluded, removal of the earplug in each implanted ferret resulted in an immediate return of sound localization accuracy in both the left and right hemifields toward pre-plug scores. **b**-**d**, Comparable recovery in spatial response properties in the IC, illustrated for the same animals by the differences between pre-plug, late plug and post-plug sessions in the azimuth RMSEs for the population-pattern decoder (**b**), the peak in the left-right decoding weights following stimulus presentation in each IC (blue: left hemifield; red: right hemifield; for clarity this is shown for F2111 only) (**c**), and the distribution of centroids and ERRF widths for all units in each implanted ferret (**d**).

## Discussion

We recorded neural activity bilaterally from the auditory midbrain as ferrets performed a sound localization task, both with normal binaural inputs and when the relationship between auditory spatial cues and sound source direction was changed by reversibly occluding one ear. The localization accuracy of neuronal populations recorded predominantly in the CNIC plus the overlying DCIC correlated with the behavioral performance of individual animals under both hearing conditions. Close correspondence with the behavior was also found when the location of the sound source was decoded from the firing rates of neurons recorded in one IC only. These findings demonstrate that spatial information encoded in the IC is sufficient to support the animals’ behavior and that plasticity in the response properties of these neurons matches the recovery of localization accuracy as ferrets adapt to conductive hearing loss in one ear.

The IC receives converging inputs from upstream brainstem nuclei that extract ITDs, ILDs and monaural spectral cues^1,39,40^, and is the first stage in the auditory pathway where integration of these cues by individual neurons takes place^3,4^. Previous measurements of spatial sensitivity under anesthesia or in awake, passively listening animals have shown that most neurons recorded in different regions of the IC have broad contralateral tuning^23,26,41–47^, with some of these studies reporting that a minority of neurons preferred ipsilateral or frontal locations^23,42,43,45,46^. Our data from behaving ferrets revealed a similar azimuthal distribution of centroid locations and that IC neurons exhibit richer temporal response patterns than those described under anesthesia in this species^48^, though sustained responses dominate in each case.

Given its pivotal role in integrating auditory spatial cues, the question of how sound source location is encoded in the IC has attracted considerable interest. In ferrets, the azimuth tuning of neurons in the nucleus of the brachium of the IC displays a coarse topographic organization^42^, which is conveyed to the SC where a more precise map of auditory space is found^49,50^. This is not the case in the CNIC in any species, however, where the presence of broad, spatial channels in each hemisphere suggests that sound azimuth may be specified by comparing the summed activity of these neuronal populations^51^. While the clearest evidence for two-channel opponent coding is found for ITD sensitivity in the CNIC^22,52,53^, other studies suggest that pattern-based models, which consider the heterogeneity of neuronal responses in the population, are more consistent with behavior, particularly in more naturalistic environments^25^ and when other spatial cues are also available^23,24^.

Previous studies have compared the neural spatial decoding performance in the IC with behavioral measurements in different animals^23,26,52^. By performing the first IC recordings during a localization task, we were able to relate neural decoding and behavioral errors in the same individuals and examine how this relationship changed during adaptation to altered cues. We found that the performance of the pattern decoder trained with activity recorded bilaterally in the IC closely matched the ferrets’ localization behavior, whereas the two-channel decoder performed less well, particularly when the animals were tested with one ear occluded. These findings therefore emphasize the importance of analyzing the responses of individual neurons when comparing azimuth tuning with behavior.

Restricting the population-pattern decoder to one IC resulted in a small but significant deficit in decoding the left-right dimension of the sound source under normal hearing conditions but no change in front-back or azimuth decoding. Other studies have also reported no differences in classifying ipsilateral and contralateral azimuths from the responses of one IC^23,24,26^. This may seem surprising given that unilateral IC lesions produce contralateral localization deficits^33–36^. In ferrets, however, significant contralateral impairments in a minimum audible angle task were found only when the lesions extended beyond the IC to include neighboring areas, such as the dorsal nucleus of the lateral lemniscus^34^. Furthermore, this lateralized behavioral deficit may reflect the loss of modulatory effects on spatial processing provided by commissural connections between each IC^54^ or the greatly reduced input to thalamocortical circuits from the lesioned side of the midbrain.

To further explore how populations of IC neurons encode sound source location and to provide insight into the plasticity of this representation, we altered the cues available for localizing sound by reversibly plugging one ear. The resulting degradation in spatial response properties and reduction in decoding accuracy are consistent with the attenuating effects of the earplug on the balance of ipsilateral inhibition and contralateral excitation to IC neurons^39,48^. Following the initial decline in localization accuracy when the left ear was plugged, the behavioral performance of all 3 implanted ferrets recovered with daily training to differing degrees. In each case, the neural decoding and behavioral errors co-varied across the sessions with the earplug in place, showing that neural correlates of adaptation are found in the IC.

Training-dependent plasticity in the IC appears to involve complementary changes in each hemisphere. In case F2108, the decoding weights of units recorded on the side contralateral to the plugged ear increased in parallel with a reduction in response inhibition and partial recovery in their spatial preferences. Recordings in anesthetized chinchillas also suggest that compensatory reduction in inhibitory input to the contralateral IC is induced by wearing an earplug in one ear for 6 weeks^55^. This is consistent with an adaptive shift in ILD sensitivity during monaural occlusion, which has been demonstrated psychophysically in adult humans^56^ and in A1 of ferrets raised with a unilateral earplug^38^.

In addition to recalibrating binaural cue sensitivity, spatial adaptation to hearing loss in one ear can be achieved through increased dependence on the unchanged monaural spectral cues provided by the open ear^27,32,56–58^. Consistent with this, we found in all 3 implanted ferrets that the decoding performance of neurons in the IC contralateral to this ear was significantly correlated with the animals’ behavior during training with the other ear occluded. Indeed, in the 2 animals with bilateral recordings, neuronal populations in this IC localized the sound as accurately as data combined from both hemispheres. The finding that individual IC neurons are capable of multiplexing multiple spatial cues using different neural encoding schemes suggests a possible basis for cue reweighting^3^.

When normal binaural cues were restored following earplug removal, the behavioral and neural responses immediately shifted back toward pre-plug values. This improvement in performance demonstrates the importance of binaural cues and supports previous studies showing that cue reweighting is rapidly reversible^29,57^. The lower decoding and behavioral performance in the first post-plug session are also consistent with evidence that earplug removal can result in a small and transient aftereffect^18,27^, indicative of plasticity in binaural sensitivity. Spatial processing in the IC therefore closely matches sound localization behavior before, during and after monaural occlusion.

Silencing A1 during stimulus presentation impairs earplug adaptation in adult ferrets but not the retrieval of previous learning when abnormal spatial cues are experienced again^18^. Because A1 corticocollicular projection neurons are essential for learning in this localization task^21^, it is likely that the plasticity observed in the IC reflects top-down cortical modulation. These inputs primarily target the dorsal and lateral cortices of the IC, with the dorsal IC being the focus of recent studies of corticocollicular modulation in mice^59–63^. In ferrets, however, corticollicular axons also innervate the CNIC^64^, where most units recorded in this study were located. Furthermore, the auditory cortex can mediate plasticity in the response properties of CNIC neurons^65,66^, potentially via intra-collicular circuits involving the IC shell^67^.

An important question is therefore whether the auditory corticocollicular projection shapes the spatial tuning of IC neurons during learning. Ferrets raised with one ear occluded develop near normal localization accuracy, which is accompanied by compensatory changes in spatial processing in A1^29,38^. The adaptive plasticity observed in the IC by training adult earplugged ferrets might therefore follow corresponding changes in A1, though recent work on the timing of cortical and subcortical plasticity argues against this^68^. Alternatively, since consolidation of learning appears to take place downstream of cortex^14,18,69^, it is possible that behavioral plasticity is driven by the changes in auditory spatial processing in the IC, which would be consistent with other evidence implicating the auditory midbrain in the control of sound-evoked movements^59,70^.

## Online Methods

A total of 11 adult female ferrets (Mustela putorius), sourced from Marshal BioResources and aged 6 months old approximately at the start of the experiments, were included in this study. All procedures were approved by the Committee on Animal Care and Ethical Review at the University of Oxford and licensed by the Home Office under the Animals (Scientific Procedures) Act (1986).

### Sound localization: hardware and training

Ferrets (n = 11) were trained by positive reinforcement to perform an approach-to-target sound localization task in a circular arena (Ø 140 cm) located inside a soundproof chamber. The arena was equipped with 12 loudspeakers (FRS 8, Visaton) for presenting target sounds, plus associated waterspouts for providing water rewards, which were distributed around the perimeter at a 30° separation. Acoustic stimuli were generated using MATLAB (MathWorks) and presented using an RX8 multi I/O processor and two SA-8 power amplifiers (Tucker-Davies Technologies). The output of each loudspeaker was digitally matched and flattened (±5 dB) across sound frequencies from 0.25-30 kHz. Training sessions were performed in blocks of decreasing broadband noise duration (2000, 1000, 500, 200, 100, 40 and 20 ms) until performance stabilized. Within each session, the sound level was randomized from 56 – 84 dB SPL in steps of 7 dB.

Ferrets were trained to stand on a platform at the center of the testing arena facing the loudspeaker at 0° location and lick a waterspout for a variable amount of time (300-500 ms) to trigger presentation of a target sound from one of the 12 loudspeakers. The animal then had to approach the location of the loudspeaker that presented the sound to receive a water reward. During periods of behavioral testing, the ferrets had access to water contingent on their performance during the twice-daily testing sessions. Only correct responses were rewarded (150–250 μl).

### Adaptation to unilateral hearing loss

Reversible unilateral conductive hearing loss was induced by inserting an earplug (E-A-R Classic 3M) into the external auditory meatus, which was shaped to fit and secured in place by filling the concha of the external ear with a silicone mould (Otoform KC, Dreve Otoplastik GmbH). Otoscopic examinations and tympanograms (Kamplex KLT25 Audiometer, P.C. Werth) were performed under Domitor sedation (medetomidine hydrochloride 0.1 mg/kg i.m., Orion Pharma) both before insertion and after removal of the earplugs to check the health status of the external and middle ear. Sedation was reversed with atipamezole hydrochloride (0.5 mg/kg s.c., Antisedan, Orion Pharma). The acoustical properties of these earplugs have been characterized previously in ferrets^29,71^ and humans^56,57^.

As in previous work^18,21,27^, we trained ferrets to adapt to an earplug using 1000-ms long broadband noise bursts as the target stimuli. Although training with noise bursts as short as 40 ms in duration allows ferrets to partially recover their sound localization accuracy^27^, the higher scores and larger changes in localization accuracy produced by the earplugs with longer stimuli make it easier to investigate the neural correlates of adaptation and to compare our data with previous studies investigating the neural circuits and acoustical cues involved in training-dependent adaptation^18,21,27,72^.

### Electrode implantation

Four of the ferrets (F2108, F2109, F2110 and F2111) from which behavioral data had been collected were implanted with Neuropixels 1.0 probes in aseptic conditions under general anesthesia induced with an intramuscular injection of Domitor (medetomidine hydrochloride, 0.022 mg/kg, Orion Pharma) and Narketan (ketamine hydrochloride, 5 mg/kg, Vetoquinol). The recordings were unsuccessful in F2109, and only behavioral data were used from this animal. Once the animals were intubated and artificially ventilated, anesthesia was then maintained with IsoFlo (isoflurane, 1–3% with 100% oxygen as a carrier, Abbott Laboratories) and the effects of Domitor reversed by administration of Antisedan (atipamezole hydrochloride, 0.06 mg/kg, s.c., Orion Pharma). Further medication administered comprised atropine (0.06 mg/kg, s.c., Hameln Pharmaceuticals) to reduce pulmonary secretions, amoxicillin/clavulanic acid (Co-amoxiclav, 20/4 mg/kg every two hours throughout the surgery, i.v., MSD Sandoz) antibiotic to prevent infections, Metacam (meloxicam, 0.2 mg/kg, s.c., Boehringer Ingelheim) and Vetergesic (buprenorphine hydrochloride, 0.03 mg/kg, s.c., Alstoe Animal Health) for analgesia, Dopram (doxapram hydrochloride, 0.2 mg/kg, s.c., Norbrook Laboratories) to prevent respiratory depression, and Dexadreson (dexamethasone, MSD Animal Health) to reduce brain edema. Heart rate, respiratory rate, oxygen saturation, and CO_2_ levels were continuously monitored. Body temperature was measured using a rectal probe and maintained at 38-39° using an air-warming BearHugger system and an electrical heating blanket. Fluids (0.9% sodium chloride, pH 7.2–7.4, plus 5% glucose) were provided at a rate of 3.0–5.0 mL/h through a cannula inserted in the radial vein. Corneal desiccation was prevented using Viscotears liquid gel (Carbomer, Bausch & Lomb).

A midline scalp incision was made and the temporal muscles retracted to expose the skull, which was cleaned ahead of performing the craniotomies. In ferrets F2108 and F2111, bilateral craniotomies were performed above the cortex overlying the IC, while in F2110 the craniotomy was made on the left side only. In two of the animals (F2110 and F2111), the exposed occipital cortex in the left craniotomy was aspirated to reveal the IC to ensure a dorsoventral probe implantation in the central nucleus. This surgery has previously been shown not to affect the ability of ferrets to localize sound or to adapt with training to unilateral conductive hearing loss^21^. Equivalent stereotaxic coordinates were then used for the right IC probe implantation for which only the dura was punctured to allow the insertion of the probe. In F2108, neural responses to acoustic stimulation were recorded during probe implantation, enabling the IC to be targeted without the need for any cerebral cortex aspiration. In this animal, anesthesia was maintained using an i.v. infusion of Domitor (0.022 mg/Kg/h) and Narketan (0.022 & 5 mg/Kg/h) in saline throughout the surgery to avoid the reduction in neural responses commonly associated with IsoFlo anesthesia.

Once the probes were in place and the craniotomies sealed with silicone (Kwik-Sil™, World precision Instruments), they were secured using bone cement (CMW1 loaded with gentamicin, DePuy) and adhesive (SuperBond, Sun Medical). The probes were electrically grounded to previously implanted bone screws, which also provided extra fixation points for the bone cement. An individually adapted rigid plastic cylindric enclosure (Ø 2.8, H 4 cm) with a narrowed base was placed around the exposed part of the probes to protect them and secured using bone cement. These enclosures had a removable cap to provide access to the probes during recording sessions. Finally, the temporal muscles were repositioned, and the scalp was trimmed and sutured around the enclosure.

The postoperative analgesic protocol consisted of 2 days of treatment with Vetergesic (0.1 mg kg/ml, s.c.) twice a day and 7 days of oral Metacam (2 mg/kg orally) and prophylactic antibiotic (amoxicillin/clavulanic acid, Synulox, 20 mg/kg, s.c.). Behavioral testing and recordings commenced approximately 2 weeks after the surgery, once the wound around the implant had healed and animals had been habituated to the handling required for connecting the probes.

### Recording frequency response areas

To further characterize the location of the recording probes within the IC, we measured the frequency tuning of units recorded under Domitor sedation by presenting 100-ms pure tones at different frequencies (499 Hz to 30 kHz) and levels (10 to 90 dB SPL in steps of 10 dB) from the loudspeaker at 0° degrees, faced by the animal when positioned on the central platform in the testing chamber. Each frequency/level combination was presented 10 times, resulting in 2700 sound presentations per recording session. Frequency response areas (FRA) were computed by summing activity in the first 50 ms window following sound onset and the resulting spike matrix was smoothed using a nine-point running hamming window^73^.

### Recordings in awake behaving animals: training and hardware

Electrophysiological data were recorded from freely moving ferrets (n=3) while they performed the approach-to-target sound localization task. Recordings were performed twice daily during the behavioral sessions, first under normal hearing conditions, then with an earplug in the left ear, to track the neural changes associated with learning-induced adaptation, and again following earplug removal.

At the beginning of any recording session, the Neuropixels probes were connected to the headstages, which were in turn connected to the control unit by an interface cable. A weight-balanced pulley system allowed the interface cable to extend and retract according to the animal’s movements in the arena without adding any tension to its head. Single and multi-unit activity was recorded from 768 recording channels simultaneously in ferrets implanted bilaterally in the IC (n = 2) or 384 channels from the single probe implanted in case F2110. The electrophysiological recordings were visualized and stored using SpikeGLX software. Digitized neural data streams from the probe headstages were registered by a Neuropixels acquisition module (PXIe, IMEC) and synchronized with the task-related events via a PXIe-6341 board (National Instruments). Thirteen digital channels (0-12) recorded the timing of the licks at the central waterspout and at each of the peripheral waterspouts, and an analog channel registered the timing of sound presentations. MATLAB (MathWorks) was used to control the trial presentation, save the trial variables, including the level and location of the presented sound, the animals’ response location and the reward given.

### Data analysis

Raw data were processed using the ecephys_spike_sorting pipeline for recordings in SpikeGLX (https://github.com/jenniferColonell/ecephys_spike_sorting.git). Raw data were filtered and denoised using CatGT. By default, as recommended by the developer due to its robustness against outlier values, a global common average referencing (gblcar) was used along with a time shift (tshift) to account for the fact that not all channels are sampled together, thereby allowing for their temporal alignment. Exceptionally, for a subgroup of sessions that contained high-frequency artefacts due to external electrical noise, the gbldmx (global demux common average referencing) filter was used instead, which performs common average referencing using groups of channels that are digitized at the same time. For sessions where noise was not present, no clear difference was observed between the two filters. Spike sorting was performed using Kilosort 3.0^74^ and was followed by a Kilosort postprocessing step which removed spike duplicates that can emerge from the sorting process. A further processing step identified noise templates based on features that conflicted with the physical properties of the unit’s action potential. Finally, neural and task events were aligned offline using Tprime, which aligns both streams using a common signal consisting of a 1 Hz square wave generated from the Imec PXIe Card.

### Unit selection for data analysis

Both single- and multi-unit responses were used for data analysis. Only units showing sound-driven responses, defined as a significant difference in firing rate within a 100 ms window after the onset or offset of a sound relative to an equivalent window prior to the presentation of the sound, were used in the analysis. Responsive units were found throughout the full length of the probes in animals F2108 and F2110. At the tip of both probes in ferret F2111, we observed a distinct response profile characterized by a periodic increase in firing, with 2-3 waves of activity and a delayed onset at ∼200 ms. This type of response differed from the short-latency onset responses typically recorded at other sites along the probes in F2111 and in other animals. These regions in F2111 were therefore excluded from further analysis. For the other animals, units recorded along the full length of the probes were analyzed.

### Analysis of response types

Single-units were classified into six main types based on their sound-evoked responses. Criteria were established for automated classification based on the activity recorded in 50-ms time windows starting at different times relative to the stimulus. Excitatory responses were defined by an increase in firing rate following stimulus onset of at least 1.5 × that measured before sound presentation. Units exhibiting excitatory responses were further classified into four groups based on differences in firing rate in each response window. If the increase in firing was confined to the onset of the stimulus, the neuron was classified as ‘onset’ type. Units that showed a persistent increase in firing rate beyond the onset of the sound were further classified into three different response profiles according to the following criteria. When the onset was followed by a decrease in activity back to pre-sound rate followed, in turn, by increased firing throughout the duration of the stimulus, the cluster was classified as ‘onset with build-up’. A decrease in activity following the onset response but which was still significantly above the pre-sound rate and maintained until the offset of the sound constituted the ‘onset sustained’ type. In some cases, this sustained component was not significantly different from the pre-sound rate but pre-sound activity was significantly higher than post-sound, so the response was still considered to be ‘onset sustained’. Finally, an increase in activity at the onset of the sound, which was maintained at a similar rate throughout the stimulus, was classified as a ‘sustained’ response. A minority of neurons showed suppressed responses to sound, characterized by a decreased firing rate following the stimulus onset relative to preceding activity. This inhibitory response could be further classified into two categories, ‘onset suppression’ and ‘sustained suppression’, based on whether the reduction in firing was limited to the sound onset or maintained throughout the stimulus, respectively.

### Spatial tuning

We found no systematic differences in the spatial response properties of single and multiunit activity, and therefore combined them in all analyses, as has been done in other studies (e.g., Lee & Middlebrooks, 2011^75^). To quantify spatial tuning, we computed the equivalent rectangular receptive field (ERRF) for each unit^75^. The ERRF is defined by the width and height of a rectangle equivalent in area to the area under the rate-azimuth function computed for the first 200 ms after sound onset. The ERRF width provides a measure of the sharpness of the azimuth tuning, and its height corresponds to the peak firing rate. The overall spatial preference was determined by the angular position of the centroid vector, calculated as the vector sum of the neural responses at each azimuthal location. Finally, to compare the overall firing rate in response to sounds presented in the left and right hemifields during the earplugged sessions, the weighted average firing rate was computed for each location (omitting the 0° and 180° positions) across all units in each session and the values for the right hemifield were subtracted from those for the left hemifield (ΔWavgFR (L – R)).

### Population linear pattern decoder

A spike-pattern decoding model was used to decode sound location from population activity in the IC. Previous studies have shown that auditory localization accuracy in ferrets, as in other species, varies with the location of the target sound^31,76^. The decline in localization accuracy observed with decreasing stimulus duration arises primarily from an increase in back-front errors, rather than left-right errors, with most of those errors occurring in the posterior hemifield^31^. Therefore, instead of treating the 12 loudspeaker locations as equivalent categories and decoding them directly, we decoded the projections of each location along the back-front and left-right axes, from which the azimuthal target location was calculated trigonometrically. This enabled us to explore in more detail how localization behavior changed after plugging one ear.

Spiking activity from each unit recorded on a probe was divided into 25 ms bins (optionally normalizing each unit’s firing rate, so that the sum of all of its spikes across trials was equal to 1), and responses were collated for each trial, giving a tensor of neuronal activity, *X_thu_*, where *t* is the trial number, *h* is the bin number within that trial, and *u* is the unit number. We also decoded target location using a collapsed two-channel model, which was similar to the pattern decoder except that all spikes from each side of the brain were collapsed into a single time-varying spike rate. We then created a target vector, *y_t_,* where *t* again corresponds to trial number. Depending on the quantity to be decoded, *y_t_* was the location in the left-right dimension of the sound source (between -1 and 1), the back-front location (between -1 and 1) or the azimuthal location of the sound source (-180 to +180 degrees). We used a linear model to produce 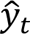, an estimate of, *y_t_*, by linearly decoding the neuronal responses:

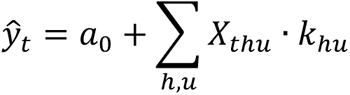

We optimized the model parameters *a*_0_ and *k_hu_* to minimize the mean square error between 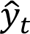 and *y_t_*, subject to L2 regularization, using glmnet (https://glmnet.stanford.edu/articles/glmnet.html). The trials were randomly shuffled and split into non-overlapping sets of 12 trials (with the last set containing any extra trials), and each of these was used as the test set for one ‘fold’ of the data. For each fold, the model was trained only on the trials that were not part of the test set for that fold. Consequently, the training and test sets were non-overlapping in every case. For each fold, the model was evaluated on the test data by measuring the mean square error (or correlation coefficient) between 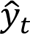 and *y_t_*. To test the models’ performance at decoding azimuth, we converted the left-right (sine of the azimuth angle) and back-front (cosine of the azimuth angle) model predictions into azimuthal angles and compared these with the actual target stimulus azimuth. For reporting accuracy in the ipsilateral and contralateral sides, root mean square errors (RMSE) were computed individually for each hemifield, including 0° and 180° in both sides.

When the decoder was trained with data recorded bilaterally in the IC, the number of neurons in each IC was equalized by randomly removing units from the side with more units. For each trial, we measured the spiking activity during the sound presentation (1000 ms in length), plus 200 ms before and after the stimulus. Additional analyses were carried out to control for the effects of head movements during the sound presentation by decoding stimulus location using data from the first 200 ms after sound onset only since this corresponds to the typical latency of sound-evoked head orienting responses in ferrets^31,32^.

The behavioral trials for each session were randomly shuffled and split into non-overlapping training (12 trials) and test sets for 24 cross-validation runs (folds). The average decoding accuracy was reported for each set.

### Histology

Once the behavioral and neural recording data had been acquired, animals were euthanized and the brain recovered to identify the location of the recording probes. Animals were euthanized under sedation (Domitor: medetomidine hydrochloride, 0.022 mg/kg, i.m., Orion Pharma) by an injection of pentobarbital sodium (Euthatal, 400 mg/kg, i.p., Merial Animal Health). Once respiration had ceased, animals were perfused transcardially with 500ml of saline (0.9% NaCl), followed by two liters of paraformaldehyde (4%) in 0.1M phosphate buffer, pH = 7.4. The brain was removed from the cranium following the direction of the probes and placed in a cryoprotective sucrose solution (30% sucrose in 0.1M PB) until it sank. The brain was blocked and the region containing the auditory midbrain was sectioned at 45-µm thickness in the coronal plane using a freezing microtome (Leica SM2000 R sliding microtome). Sections containing the IC were mounted on gelatinized slides and stained for Nissl substance. Finally, slides were cover-slipped using mounting medium (Entellan, Merck).

Anatomical subdivisions of the IC were identified in each section and drawn under the microscope (DMR Leica Microsystems) fitted with a camera lucida. IC subdivisions were identified based on previous descriptions of cell morphology and density in different species, including the ferret^64,77^. The location of the probes was readily apparent under the microscope from the physical lesion left in the tissue and the associated gliosis, characterized by much denser Nissl staining of small cells at the tip or along the track of the probe. Examination of the serial sections allowed reconstruction of probe track in the IC when its trajectory did not match the plane of sectioning.

### Statistical analysis

To directly compare the accuracy of each ferret’s localization behavior with the performance of the IC population decoders for that animal, we computed the RMSE for each measure and fitted regression lines and their 95% confidence intervals (shaded areas). Linear regression was also used to measure the rate of training-induced adaptation in the behavioral responses and the rate of any accompanying change in IC decoding accuracy. Normalized firing rates, ERRF width, and centroids were compared between normal hearing and earplugged conditions using different statistical tests according to the underlying distribution of the data, as stated in each case. All statistical analyses were performed using RStudio (RStudio: Integrated Development for R. RStudio, Inc., Boston, MA, United).

## Notes

### Competing Interest Statement

The authors have declared no competing interest.

### Funding

- Wellcome Trust: WT108369
- Royal National Institute for Deaf People: S55_King

## References

1 Grothe, B., Pecka, M. & McAlpine, D. Mechanisms of sound localization in mammals. Physiol Rev 90, 983–1012 (2010). 10.1152/physrev.00026.2009

2 Grothe, B. & Pecka, M. The natural history of sound localization in mammals--a story of neuronal inhibition. Front Neural Circuits 8, 116 (2014). 10.3389/fncir.2014.00116

3 Chase, S. M. & Young, E. D. Cues for sound localization are encoded in multiple aspects of spike trains in the inferior colliculus. J Neurophysiol 99, 1672–1682 (2008). 10.1152/jn.00644.2007

4 Chase, S. M. & Young, E. D. Limited segregation of different types of sound localization information among classes of units in the inferior colliculus. J Neurosci 25, 7575–7585 (2005). 10.1523/jneurosci.0915-05.2005

5 Anderson, L. A., Malmierca, M. S., Wallace, M. N. & Palmer, A. R. Evidence for a direct, short latency projection from the dorsal cochlear nucleus to the auditory thalamus in the guinea pig. Eur J Neurosci 24, 491–498 (2006). 10.1111/j.1460-9568.2006.04930.x

6 Schofield, B. R., Mellott, J. G. & Motts, S. D. Subcollicular projections to the auditory thalamus and collateral projections to the inferior colliculus. Front Neuroanat 8, 70 (2014). 10.3389/fnana.2014.00070

7 Ito, T., Bishop, D. C. & Oliver, D. L. Functional organization of the local circuit in the inferior colliculus. Anat Sci Int 91, 22–34 (2016). 10.1007/s12565-015-0308-8

8 King, A. J. Sensory experience and the formation of a computational map of auditory space in the brain. Bioessays 21, 900–911 (1999). 10.1002/(sici)1521-1878(199911)21:11<900::Aid-bies2>3.0.Co;2-6

9 Keating, P. & King, A. J. Sound localization in a changing world. Curr Opin Neurobiol 35, 35–43 (2015). 10.1016/j.conb.2015.06.005

10 Keating, P. & King, A. J. Developmental plasticity of spatial hearing following asymmetric hearing loss: context-dependent cue integration and its clinical implications. Front Syst Neurosci 7, 123 (2013). 10.3389/fnsys.2013.00123

11 Kumpik, D. P. & King, A. J. A review of the effects of unilateral hearing loss on spatial hearing. Hear Res 372, 17–28 (2019). 10.1016/j.heares.2018.08.003

12 Mendonca, C. A review on auditory space adaptations to altered head-related cues. Front Neurosci 8, 219 (2014). 10.3389/fnins.2014.00219

13 Caras, M. L. & Sanes, D. H. Top-down modulation of sensory cortex gates perceptual learning. Proc Natl Acad Sci U S A 114, 9972–9977 (2017). 10.1073/pnas.1712305114

14 Drieu, C. et al. Rapid emergence of latent knowledge in the sensory cortex drives learning. Nature 641, 960–970 (2025). 10.1038/s41586-025-08730-8

15 Maor, I. et al. Neural correlates of learning pure tones or natural sounds in the auditory cortex. Front Neural Circuits 13, 82 (2020). 10.3389/fncir.2019.00082

16 Polley, D. B., Steinberg, E. E. & Merzenich, M. M. Perceptual learning directs auditory cortical map reorganization through top-down influences. Journal of neuroscience 26, 4970–4982 (2006).

17 Rutkowski, R. G. & Weinberger, N. M. Encoding of learned importance of sound by magnitude of representational area in primary auditory cortex. Proc Natl Acad Sci U S A 102, 13664–13669 (2005). 10.1073/pnas.0506838102

18 Bajo, V. M. et al. Silencing cortical activity during sound-localization training impairs auditory perceptual learning. Nat Commun 10, 3075 (2019). 10.1038/s41467-019-10770-4

19 Nodal, F. R., Bajo, V. M. & King, A. J. Plasticity of spatial hearing: behavioural effects of cortical inactivation. J Physiol 590, 3965–3986 (2012). 10.1113/jphysiol.2011.222828

20 Trapeau, R. & Schonwiesner, M. Adaptation to shifted interaural time differences changes encoding of sound location in human auditory cortex. Neuroimage 118, 26–38 (2015). 10.1016/j.neuroimage.2015.06.006

21 Bajo, V. M., Nodal, F. R., Moore, D. R. & King, A. J. The descending corticocollicular pathway mediates learning-induced auditory plasticity. Nat Neurosci 13, 253–260 (2010). 10.1038/nn.2466

22 McAlpine, D., Jiang, D. & Palmer, A. R. A neural code for low-frequency sound localization in mammals. Nat Neurosci 4, 396–401 (2001). 10.1038/86049

23 Day, M. L. & Delgutte, B. Decoding sound source location and separation using neural population activity patterns. J Neurosci 33, 15837–15847 (2013). 10.1523/JNEUROSCI.2034-13.2013

24 Day, M. L. & Delgutte, B. Neural population encoding and decoding of sound source location across sound level in the rabbit inferior colliculus. J Neurophysiol 115, 193–207 (2016). 10.1152/jn.00643.2015

25 Goodman, D. F., Benichoux, V. & Brette, R. Decoding neural responses to temporal cues for sound localization. Elife 2, e01312 (2013). 10.7554/eLife.01312

26 Boffi, J. C., Bathellier, B., Asari, H. & Prevedel, R. Noisy neuronal populations effectively encode sound localization in the dorsal inferior colliculus of awake mice. Elife 13 (2024). 10.7554/eLife.97598

27 Kacelnik, O., Nodal, F. R., Parsons, C. H. & King, A. J. Training-induced plasticity of auditory localization in adult mammals. PLoS Biol 4, e71 (2006). 10.1371/journal.pbio.0040071

28 Eric Lupo, J., Koka, K., Thornton, J. L. & Tollin, D. J. The effects of experimentally induced conductive hearing loss on spectral and temporal aspects of sound transmission through the ear. Hear Res 272, 30–41 (2011). 10.1016/j.heares.2010.11.003

29 Keating, P., Dahmen, J. C. & King, A. J. Context-specific reweighting of auditory spatial cues following altered experience during development. Curr Biol 23, 1291–1299 (2013). 10.1016/j.cub.2013.05.045

30 Moore, D. R., Semple, M. N. & Addison, P. D. Some acoustic properties of neurones in the ferret inferior colliculus. Brain Res 269, 69–82 (1983). 10.1016/0006-8993(83)90963-0

31 Nodal, F. R., Bajo, V. M., Parsons, C. H., Schnupp, J. W. & King, A. J. Sound localization behavior in ferrets: comparison of acoustic orientation and approach-to-target responses. Neuroscience 154, 397–408 (2008). 10.1016/j.neuroscience.2007.12.022

32 Sanchez Jimenez, A., Willard, K. J., Bajo, V. M., King, A. J. & Nodal, F. R. Persistence and generalization of adaptive changes in auditory localization behavior following unilateral conductive hearing loss. Front Neurosci 17, 1067937 (2023). 10.3389/fnins.2023.1067937

33 Jenkins, W. M. & Masterton, R. B. Sound localization: effects of unilateral lesions in central auditory system. J Neurophysiol 47, 987–1016 (1982). 10.1152/jn.1982.47.6.987

34 Kelly, J. B. & Kavanagh, G. L. Sound localization after unilateral lesions of inferior colliculus in the ferret (*Mustela putorius*). J Neurophysiol 71, 1078–1087 (1994). 10.1152/jn.1994.71.3.1078

35 Champoux, F. et al. Auditory processing in a patient with a unilateral lesion of the inferior colliculus. Eur J Neurosci 25, 291–297 (2007). 10.1111/j.1460-9568.2006.05260.x

36 Litovsky, R. Y., Fligor, B. J. & Tramo, M. J. Functional role of the human inferior colliculus in binaural hearing. Hear Res 165, 177–188 (2002). 10.1016/s0378-5955(02)00304-0

37 Stecker, G. C., Harrington, I. A. & Middlebrooks, J. C. Location coding by opponent neural populations in the auditory cortex. PLoS Biol 3, e78 (2005). 10.1371/journal.pbio.0030078

38 Keating, P., Dahmen, J. C. & King, A. J. Complementary adaptive processes contribute to the developmental plasticity of spatial hearing. Nat Neurosci 18, 185–187 (2015). 10.1038/nn.3914

39 Pollak, G. D. Circuits for processing dynamic interaural intensity disparities in the inferior colliculus. Hear Res 288, 47–57 (2012). 10.1016/j.heares.2012.01.011

40 Ryugo, D. K. & Milinkeviciute, G. Differential projections from the cochlear nucleus to the inferior colliculus in the mouse. Front Neural Circuits 17, 1229746 (2023). 10.3389/fncir.2023.1229746

41 Aitkin, L. M., Gates, G. R. & Phillips, S. C. Responses of neurons in inferior colliculus to variations in sound-source azimuth. J Neurophysiol 52, 1–17 (1984). 10.1152/jn.1984.52.1.1

42 Schnupp, J. W. & King, A. J. Coding for auditory space in the nucleus of the brachium of the inferior colliculus in the ferret. J Neurophysiol 78, 2717–2731 (1997). 10.1152/jn.1997.78.5.2717

43 Behrend, O., Dickson, B., Clarke, E., Jin, C. & Carlile, S. Neural responses to free field and virtual acoustic stimulation in the inferior colliculus of the guinea pig. J Neurophysiol 92, 3014–3029 (2004). 10.1152/jn.00402.2004

44 Groh, J. M., Kelly, K. A. & Underhill, A. M. A monotonic code for sound azimuth in primate inferior colliculus. J Cogn Neurosci 15, 1217–1231 (2003). 10.1162/089892903322598166

45 Kuwada, S., Bishop, B., Alex, C., Condit, D. W. & Kim, D. O. Spatial tuning to sound-source azimuth in the inferior colliculus of unanesthetized rabbit. J Neurophysiol 106, 2698–2708 (2011). 10.1152/jn.00532.2011

46 van den Wildenberg, M. F. & Bremen, P. Heterogeneous spatial tuning in the auditory pathway of the Mongolian Gerbil (Meriones unguiculatus). Eur J Neurosci 60, 4954–4981 (2024). 10.1111/ejn.16472

47 Yao, J. D., Bremen, P. & Middlebrooks, J. C. Transformation of spatial sensitivity along the ascending auditory pathway. J Neurophysiol 113, 3098–3111 (2015). 10.1152/jn.01029.2014

48 McAlpine, D., Martin, R. L., Mossop, J. E. & Moore, D. R. Response properties of neurons in the inferior colliculus of the monaurally deafened ferret to acoustic stimulation of the intact ear. J Neurophysiol 78, 767–779 (1997). 10.1152/jn.1997.78.2.767

49 King, A. J. & Hutchings, M. E. Spatial response properties of acoustically responsive neurons in the superior colliculus of the ferret: a map of auditory space. J Neurophysiol 57, 596–624 (1987). 10.1152/jn.1987.57.2.596

50 King, A. J., Schnupp, J. W. & Thompson, I. D. Signals from the superficial layers of the superior colliculus enable the development of the auditory space map in the deeper layers. J Neurosci 18, 9394–9408 (1998). 10.1523/jneurosci.18-22-09394.1998

51 McAlpine, D. Creating a sense of auditory space. J Physiol 566, 21–28 (2005). 10.1113/jphysiol.2005.083113

52 Belliveau, L. A., Lyamzin, D. R. & Lesica, N. A. The neural representation of interaural time differences in gerbils is transformed from midbrain to cortex. J Neurosci 34, 16796–16808 (2014). 10.1523/jneurosci.2432-14.2014

53 Lesica, N. A., Lingner, A. & Grothe, B. Population coding of interaural time differences in gerbils and barn owls. J Neurosci 30, 11696–11702 (2010). 10.1523/jneurosci.0846-10.2010

54 Orton, L. D., Papasavvas, C. A. & Rees, A. Commissural gain control enhances the midbrain representation of sound location. J Neurosci 36, 4470–4481 (2016). 10.1523/jneurosci.3012-15.2016

55 Thornton, J. L., Anbuhl, K. L. & Tollin, D. J. Temporary unilateral hearing loss impairs spatial auditory information processing in neurons in the central auditory system. Front Neurosci 15, 721922 (2021). 10.3389/fnins.2021.721922

56 Keating, P., Rosenior-Patten, O., Dahmen, J. C., Bell, O. & King, A. J. Behavioral training promotes multiple adaptive processes following acute hearing loss. Elife 5, e12264 (2016). 10.7554/eLife.12264

57 Kumpik, D. P., Kacelnik, O. & King, A. J. Adaptive reweighting of auditory localization cues in response to chronic unilateral earplugging in humans. J Neurosci 30, 4883–4894 (2010). 10.1523/JNEUROSCI.5488-09.2010

58 Zonooz, B. & Van Opstal, A. J. Differential adaptation in azimuth and elevation to acute monaural spatial hearing after training with visual feedback. eNeuro 6 (2019). 10.1523/eneuro.0219-19.2019

59 Xiong, X. R. et al. Auditory cortex controls sound-driven innate defense behaviour through corticofugal projections to inferior colliculus. Nat Commun 6, 7224 (2015). 10.1038/ncomms8224

60 Blackwell, J. M., Lesicko, A. M., Rao, W., De Biasi, M. & Geffen, M. N. Auditory cortex shapes sound responses in the inferior colliculus. Elife 9 (2020). 10.7554/eLife.51890

61 Lee, T. Y., Weissenberger, Y., King, A. J. & Dahmen, J. C. Midbrain encodes sound detection behavior without auditory cortex. Elife 12 (2024). 10.7554/eLife.89950

62 Oberle, H. M., Ford, A. N., Dileepkumar, D., Czarny, J. & Apostolides, P. F. Synaptic mechanisms of top-down control in the non-lemniscal inferior colliculus. Elife 10 (2022). 10.7554/eLife.72730

63 Weible, A. P., Yavorska, I. & Wehr, M. A cortico-collicular amplification mechanism for gap detection. Cereb Cortex 30, 3590–3607 (2020). 10.1093/cercor/bhz328

64 Bajo, V. M., Nodal, F. R., Bizley, J. K., Moore, D. R. & King, A. J. The ferret auditory cortex: descending projections to the inferior colliculus. Cereb Cortex 17, 475–491 (2007). 10.1093/cercor/bhj164

65 Gao, E. & Suga, N. Experience-dependent corticofugal adjustment of midbrain frequency map in bat auditory system. Proc Natl Acad Sci U S A 95, 12663–12670 (1998). 10.1073/pnas.95.21.12663

66 Zhang, Y., Hakes, J. J., Bonfield, S. P. & Yan, J. Corticofugal feedback for auditory midbrain plasticity elicited by tones and electrical stimulation of basal forebrain in mice. Eur J Neurosci 22, 871–879 (2005). 10.1111/j.1460-9568.2005.04276.x

67 Jen, P. H., Sun, X. & Chen, Q. C. An electrophysiological study of neural pathways for corticofugally inhibited neurons in the central nucleus of the inferior colliculus of the big brown bat, Eptesicus fuscus. Exp Brain Res 137, 292–302 (2001). 10.1007/s002210000637

68 Ying, R., Stolzberg, D. J. & Caras, M. L. Neural correlates of perceptual plasticity in the auditory midbrain and thalamus. J Neurosci 45 (2025). 10.1523/jneurosci.0691-24.2024

69 Kawai, R. et al. Motor cortex is required for learning but not for executing a motor skill. Neuron 86, 800–812 (2015). 10.1016/j.neuron.2015.03.024

70 Clayton, K. K. et al. Sound elicits stereotyped facial movements that provide a sensitive index of hearing abilities in mice. Curr Biol 34, 1605–1620.e1605 (2024). 10.1016/j.cub.2024.02.057

71 Moore, D. R., Hutchings, M. E., King, A. J. & Kowalchuk, N. E. Auditory brain stem of the ferret: some effects of rearing with a unilateral ear plug on the cochlea, cochlear nucleus, and projections to the inferior colliculus. J Neurosci 9, 1213–1222 (1989). 10.1523/jneurosci.09-04-01213.1989

72 Nodal, F. R. et al. Lesions of the auditory cortex impair azimuthal sound localization and its recalibration in ferrets. J Neurophysiol 103, 1209–1225 (2010). 10.1152/jn.00991.2009

73 Bizley, J. K., Nodal, F. R., Nelken, I. & King, A. J. Functional organization of ferret auditory cortex. Cereb Cortex 15, 1637–1653 (2005). 10.1093/cercor/bhi042

74 Pachitariu, M., Sridhar, S., Pennington, J. & Stringer, C. Spike sorting with Kilosort4. Nat Methods 21, 914–921 (2024). 10.1038/s41592-024-02232-7

75 Lee, C. C. & Middlebrooks, J. C. Auditory cortex spatial sensitivity sharpens during task performance. Nat Neurosci 14, 108–114 (2011). 10.1038/nn.2713

76 King, A. J. & Parsons, C. H. Improved auditory spatial acuity in visually deprived ferrets. Eur J Neurosci 11, 3945–3956 (1999). 10.1046/j.1460-9568.1999.00821.x

77. Radtke-Schuller, S. Cyto- and Myeloarchitectural Brain Atlas of the Ferret (Mustela putorius) in MRI Aided Stereotaxic Coordinates. (Springer Cham, 2018).

