## Extended Figures for "Auditory midbrain encodes training-induced plasticity in sound localization behavior"

Extended Figures 1-7

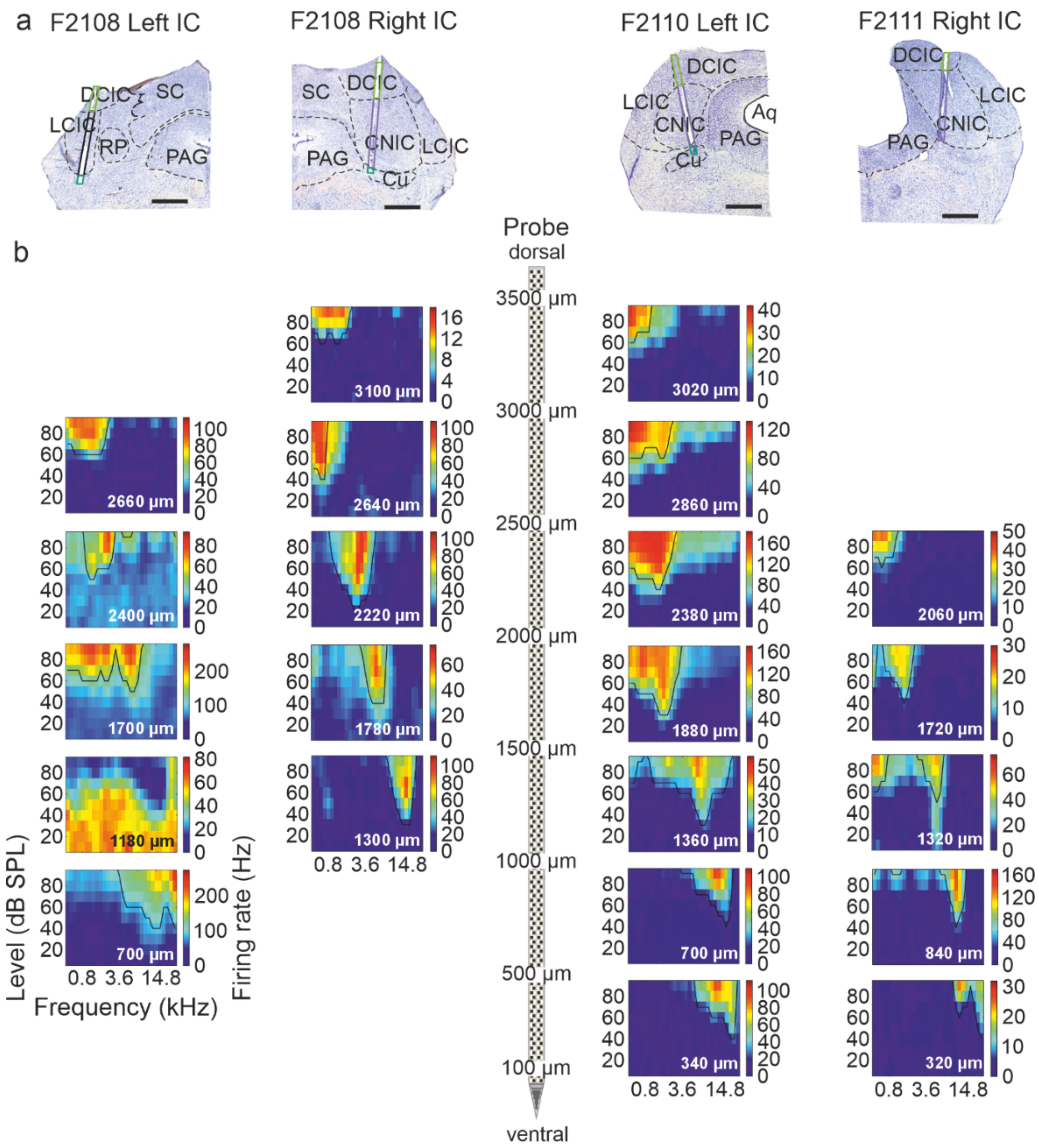


**Extended Fig. 1: Frequency response areas across recording probes reveal IC tonotopy in all animals.**

**a**, Coronal sections stained for Nissl substance at the level of the IC for ferrets F2108, F2110 and F2111, with the chronically implanted Neuropixels probes superimposed on the electrode track and color coded according to their anatomical locations. Aq, Aqueduct; DCIC, dorsal cortex of the IC; Cu, cuneiform nucleus; CNIC, central nucleus of the IC; LCIC, lateral cortex of the IC; Scale bar, 1 mm. **b**, Representative frequency response areas recorded under sedation aligned with their recording depths on the probe. Each column of frequency response areas comes from the IC shown in **a**.

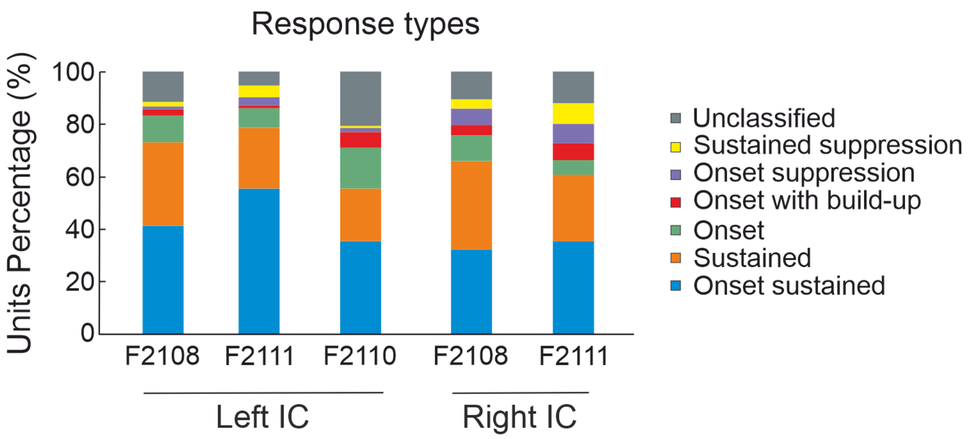


**Extended Fig. 2: Distribution of temporal firing patterns for single units recorded in the IC of each ferret.**

Histogram showing the percentage of the different response types across the different animals. Despite small variations across cases, the majority of units exhibited an excitatory response, and the most common temporal response pattern was a sustained response, sometimes accompanied by an onset component.

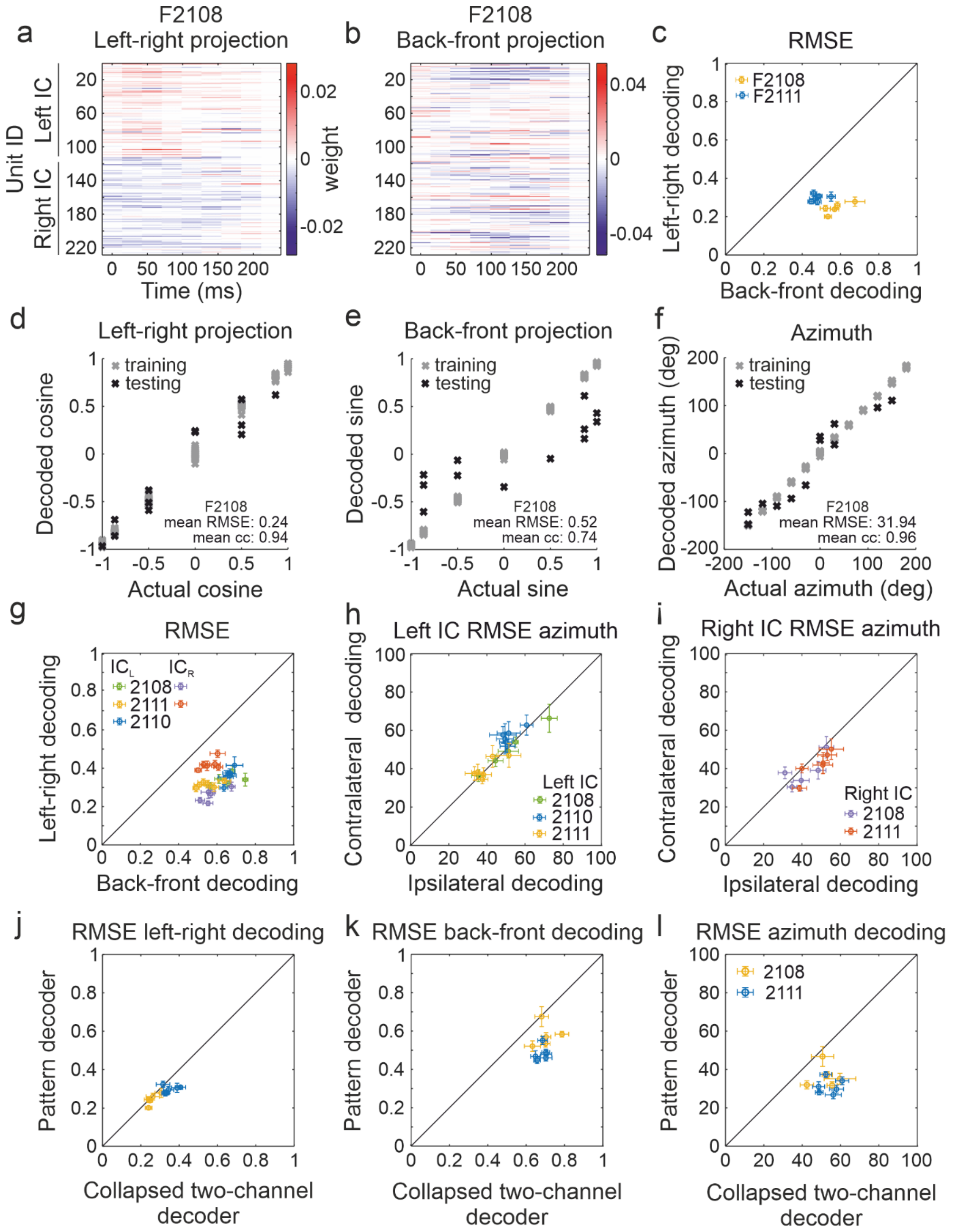


**Extended Fig. 3: A simple linear pattern model captures spatial information encoded in the IC even when restricted to the first 200 ms following target sound onset.**

The 12 target locations used in the sound localization task were decomposed into their left-right and back-front projections, which were then decoded using a linear population-pattern model from the responses recorded in the first 200 ms following target sound onset. For each session, the number of units recorded in each IC was balanced, and the model was independently fitted multiple times (*k*-folds). For each fold, 12 different trials were selected for testing the model and training was performed with the remaining trials from that session. **a**, **b**, Temporal distribution over the first 200 ms following sound onset of the unit weights for an example fold of the model (case F2108) for the left-right projection (**a**) and the back-front projection (**b**). Units are ordered according to their IC location in each hemisphere, and the color indicates how informative each unit is about the decoded variable (blue: left hemifield (**a**), back (i.e., posterior hemifield, **b**); red: right hemifield (**a**), front (anterior hemifield, **b**)). **c**, Root mean squared errors (RMSEs) for decoding the left-right vs the back-front axis for animals with bilateral IC recordings. **d**, **e**, **f**, Distribution of values for the training trials (gray) and testing trials (black) for decoding the left-right projection (cosine), back-front projection (sine), and azimuthal location (degrees), respectively, for an example fold. Overall performance across folds is expressed as the mean RMSE and mean correlation coefficient. **g**, **h**, **i**, Decoding performance based on only one IC. **g**, RMSEs for decoding the left-right vs the back-front axis for either the left or right IC. **h**, **i**, Comparison of RMSEs for decoding contralateral vs ipsilateral locations for models using only left (**h**) or right IC units (**i**). **j**, **k**, **l**, Comparison of RMSEs between the population-pattern and collapsed two-channel decoders for the left-right axis (**j**), back-front axis (**k**) and azimuthal location (**l**).

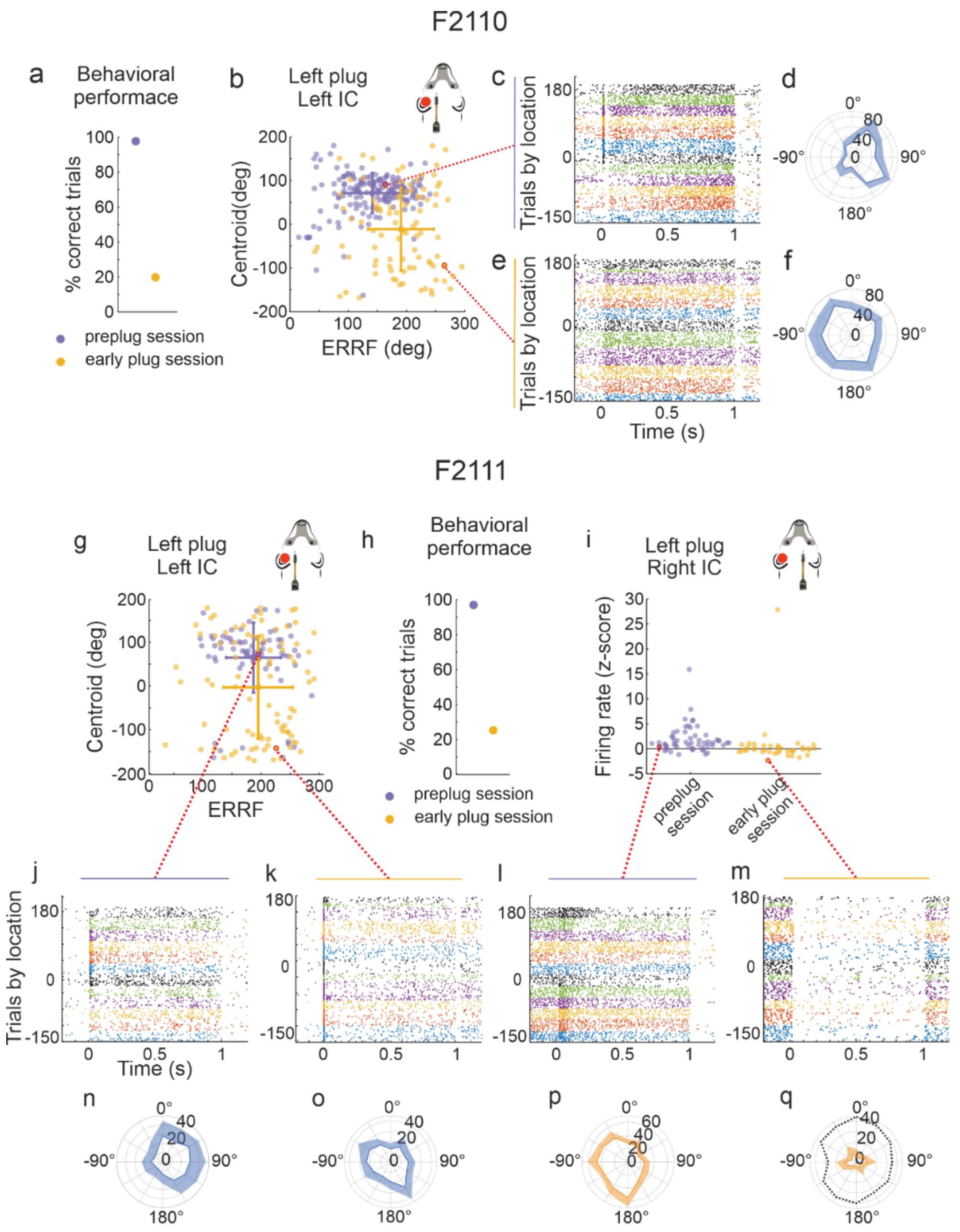


**Extended Fig. 4: Effects of monaural hearing loss on IC activity and spatial properties in cases F2110 and F2111.**

**a**, Initial reduction in sound localization accuracy caused by monaural hearing loss in case F2110. **b**, Changes in spatial response properties (centroid and ERRF width) of all units recorded in the left IC of F2110 following earplug insertion. **c,** Raster plots of example unit in the left IC before earplug insertion. **d**, Azimuth-response profile of this unit. **e,** Raster plots of an example unit in the left IC soon after earplug insertion. **f**, Azimuth-response profile of this unit. **g**, Changes in spatial response properties (centroid and ERRF width) of all units recorded in the left IC of F2111 following earplug insertion. **h**, Initial reduction in localization accuracy caused by monaural hearing loss in F2111. **i**, Suppression of sound-driven activity in the right IC of F2111 following earplug insertion. Raster plots of example units the left IC (**j**, **k**) and right IC (**l**, **m**) before and after earplug insertion. **n-q**, Azimuth response profiles of these same units. The unit in **m** and **q** was inhibited by sound and its baseline activity is shown by the dotted line in **q**. In all cases, an early plug session means in the first 1-2 days following earplug insertion.

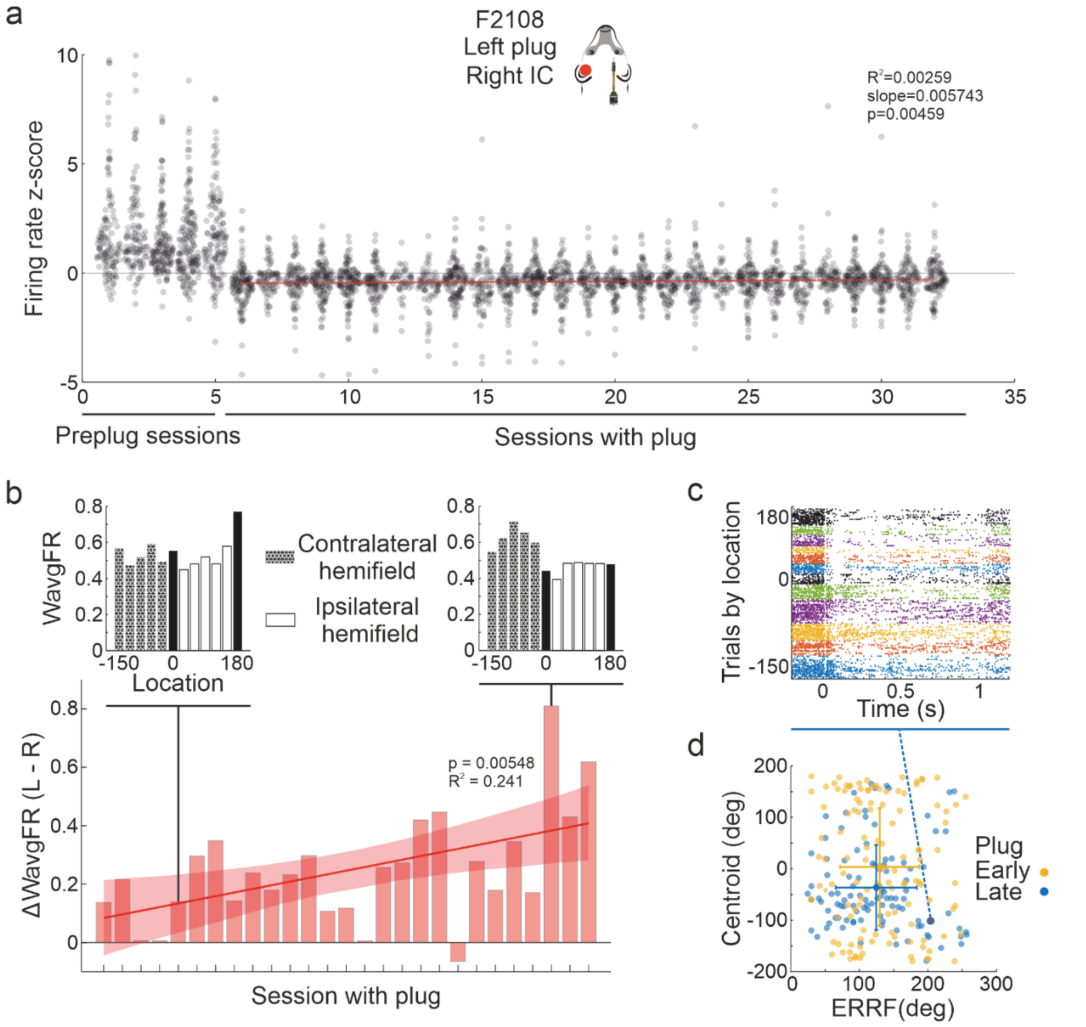


**Extended Fig. 5: Neural changes in the IC contralateral to the plugged ear over the course of sound localization training.**

**a**, Plugging the left ear resulted in inhibition of activity in the contralateral right IC, which showed a modest recovery over the period of monaural occlusion, as indicated by a significant positive slope in the regression between the firing rate z-score and number of days of wearing the earplug. Data from case F2108. **b**, A significant difference emerged over the course of monaural occlusion in the weighted average firing rate in the right IC in response to sounds presented in the left vs the right hemifields (ΔWavgFR (L - R)), indicating a partial recovery of the contralateral preference of these units (0° and 180° loudspeaker positions were excluded for this analysis). The WavgFR (L – R) is shown for each target location across all units for example early and late sessions. **c**, Raster plot for an example unit from a late plug session, showing a greater response to left hemifield sounds (negative azimuths) despite the presence of an earplug in the left ear. **d**, Distribution of centroids and ERRF widths for all units in early and late plugging sessions (same sessions as in **b**), showing a recovery over time in their contralateral preference.

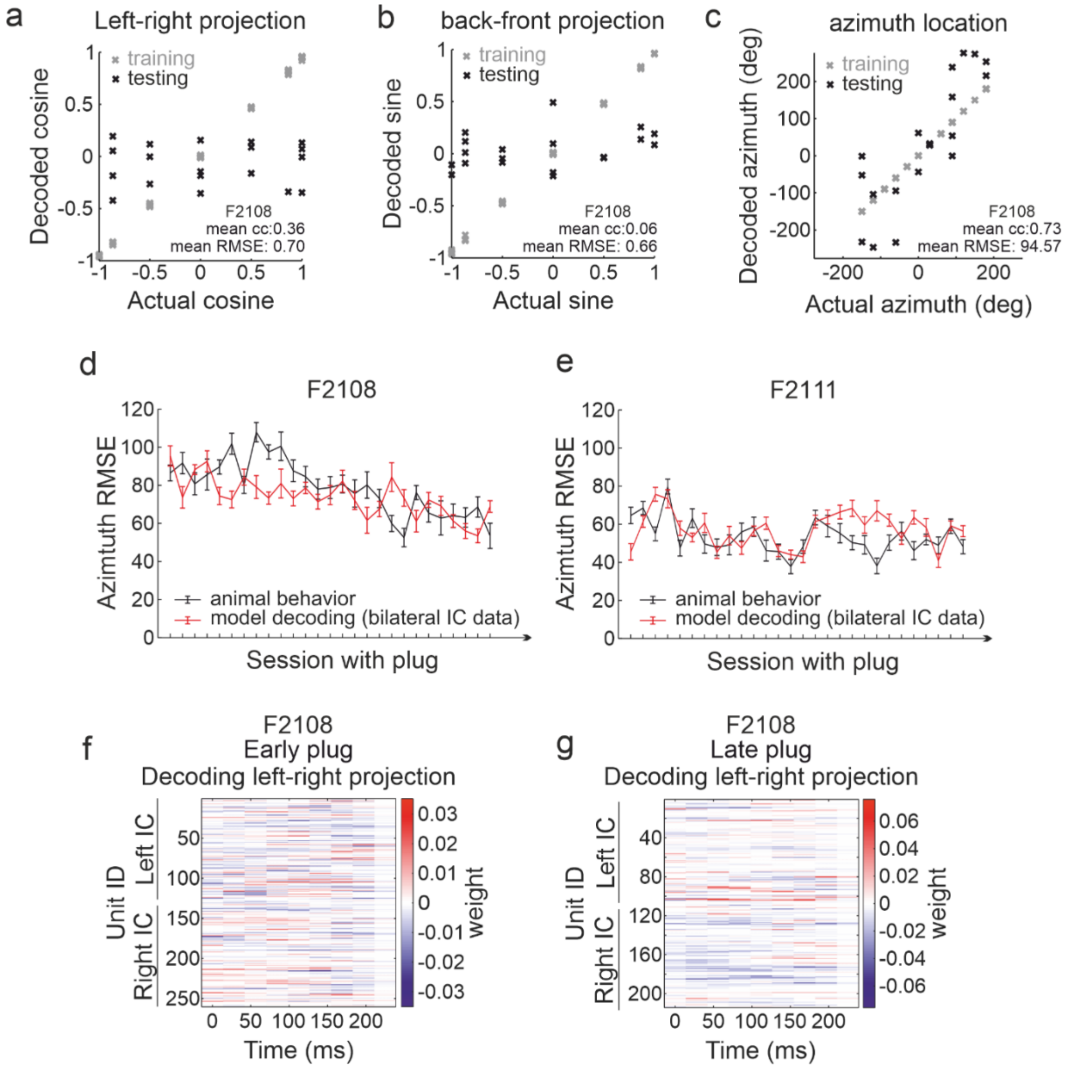


**Extended Fig. 6: Training-dependent adaptation in sound localization behavior is captured by decoding spiking activity in the inferior colliculus even when restricted to the first 200 ms following target sound onset.**

**a**-**c**, Distribution of values for the training trials (gray) and testing trials (black) for decoding the left-right axis (**a**), back-front axis (**b**), and azimuthal location (**c**) by the population-pattern model from the first 200 ms of IC activity following target sound onset for an example fold at the start of the earplugging run in case F2108. Overall performance across folds is expressed as the mean RMSE and mean correlation coefficient. **d**, **e**, Behavioral (black) and neural decoding (red) errors across the earplugging sessions for the animals with bilateral IC recordings (**d**, F2108; **e**, F2111). Only the first 200 ms of neural activity was used for decoding. **f**, **g**, Temporal distribution (over the first 200 ms following sound onset) of the unit weights for an example fold of the population-pattern model (F2108) for decoding the left-right axis in an early (**f**) and late (**g**) earplugging session. Units are ordered according to their IC location in each hemisphere, and the color indicates how informative each unit is about the decoded variable (blue: left hemifield; red: right hemifield).

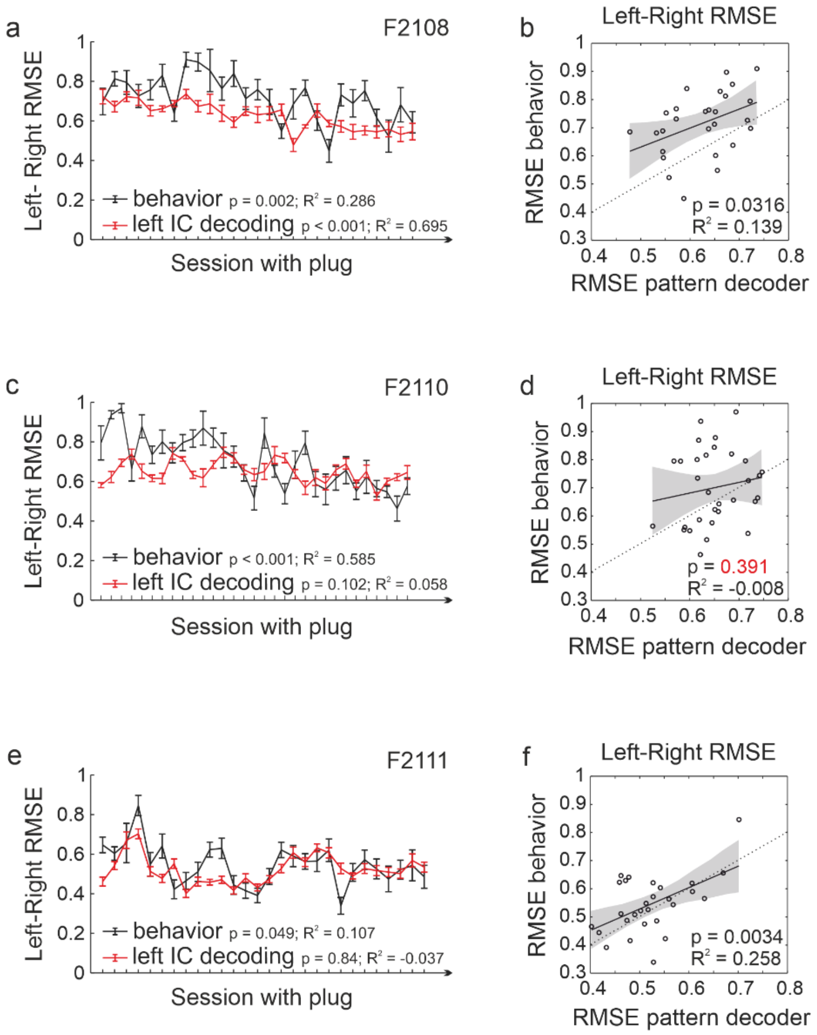


**Extended Fig. 7: Decoding localization performance from the first 200 ms of activity in one IC during ear plugging.**

**a**, **b**, Comparison of behavioral and left-right decoding errors during monaural hearing loss using activity recorded in the left IC activity for the first 200 ms following sound onset (case F2108). Spatial information was decoded from the IC using the population-pattern model. **c**, **d**, Same for F2110. **e**, **f**, Same for F2111. **a**, **c**, **e**, Covariation of behavioral (black) and IC decoder (red) RMSEs across all earplugged sessions. **b**, **d**, **f**, Linear regression between the behavioral and decoder RMSEs in each session and animal.
